# Heart Rate Variability Covaries with Amygdala Functional Connectivity During Voluntary Emotion Regulation

**DOI:** 10.1101/2022.05.06.490895

**Authors:** Emma Tupitsa, Ifeoma Egbuniwe, William K. Lloyd, Marta Puertollano, Birthe Macdonald, Karin Joanknecht, Michiko Sakaki, Carien M. van Reekum

## Abstract

The Neurovisceral Integration Model posits that shared neural networks support the effective regulation of emotions and heart rate, with heart rate variability (HRV) serving as an objective, peripheral index of prefrontal inhibitory control. Prior neuroimaging studies have predominantly examined both HRV and associated neural functional connectivity at rest, as opposed to contexts that require active emotion regulation. The present study sought to extend upon previous resting-state functional connectivity findings, examining HRV and corresponding amygdala functional connectivity during a cognitive reappraisal task. Seventy adults (52 old and 18 young adults, 18-84 years, 51% male) received instructions to cognitively reappraise negative and neutral affective images during functional MRI scanning. HRV measures were derived from a finger pulse signal throughout the scan. During the task, young adults exhibited a significant inverse association between HRV and amygdala-medial prefrontal cortex (mPFC) functional connectivity, in which higher HRV was correlated with weaker amygdala-mPFC coupling, whereas old adults displayed a slight positive, albeit non-significant correlation. Furthermore, voxelwise whole-brain functional connectivity analyses showed that higher HRV was linked to weaker right amygdala-posterior cingulate cortex connectivity across old and young adults, and in old adults, higher HRV positively correlated with stronger right amygdala – right ventrolateral prefrontal cortex connectivity. Collectively, these findings highlight the importance of assessing HRV and neural functional connectivity during active regulatory contexts to further identify neural concomitants of HRV and adaptive emotion regulation.

## 1. Introduction

The ability to flexibly respond to ongoing and complex changes in our environment, in both a timely and contextually appropriate manner, is crucial for successful adaptation to environmental challenges and emotion regulation (Aldao et al., 2015; Thompson, 1994). Responses to such situational demands generates a cascade of changes at both subjective (e.g., emotional states, expressions) and physiological (e.g., elevations or reductions to heart rate, sweating, heightened neural responding) levels. Heart Rate Variability (HRV), physiologically defined as the variation in time intervals between consecutive heart beats, has increasingly been employed as an objective, peripheral measure to capture individual differences in adaptive autonomic responding and self-regulatory capacity, including emotion regulation (Appelhans & Luecken, 2006).

HRV reflects the predominance of the parasympathetic branch of the autonomic nervous system (ANS). Both the sympathetic and parasympathetic branches directly innervate the heart via the stellate ganglia and vagus nerve respectively (Berntson et al., 1997). Dynamic interplay between both branches produces complex variations in the heart rate period that is captured by HRV, but it is the fast, modulatory impact of the parasympathetic nervous system (via the vagus nerve) that reportedly exhibits the strongest influence on the heart’s pace maker (i.e., sinoatrial node) and subsequent variation in heart rate, particularly at rest (Berntson et al., 1997). Typically, the higher the HRV, the more adaptive and responsive the cardiovascular system is, supporting fast and flexible alterations in physiological responses to effectively manage stressors, as well as maintaining homeostasis (Shaffer & Ginsberg, 2017; but see Kogan et al., 2013 for discussion on the quadratic nature of HRV).

Several models discuss the role of HRV in adaptive psychophysiological responding (Grossman & Taylor, 2007; Laborde et al., 2018; Porges, 2007, 2011; Smith et al., 2017; Thayer & Lane, 2000, 2009). In particular, the Neurovisceral Integration Model (NIM; Smith et al., 2017; Thayer & Lane, 2000, 2009) outlines a complex and reciprocal network of neural regions that overlap to support autonomic, cognitive and affective regulatory processes. At the heart of the NIM is the ‘central autonomic network’ (CAN; Benarroch, 1993), which encompasses higher cortical structures (ventromedial prefrontal cortex, anterior cingulate cortex), subcortical limbic regions (central nucleus of the amygdala, hypothalamus) and brainstem structures (periaqueductal gray, parabrachial nucleus), forming a vital, coordinated network that facilitates autonomic function and regulation (Benarroch, 1993; Thayer et al., 2009a). The NIM posits that the prefrontal cortex exerts tonic inhibitory control over subcortical structures, and by extension the vagus nerve. As such, resting HRV is proposed to serve as an index of the effective functioning of inhibitory cortical-subcortical connectivity and CNS-ANS integration, in turn promoting adaptive self-regulation (Thayer & Lane, 2000, 2009; Thayer et al., 2009a).

A growing body of neuroimaging research lends support for the NIM and the link between HRV and emotion regulation-related brain function (Mather & Thayer, 2018; Sakaki et al., 2016; Schumann et al., 2021a; Steinfurth et al., 2018). Consistent with the notion that HRV serves as a measure of effective, inhibitory cortical-subcortical connectivity, individuals with higher HRV exhibit stronger resting medial prefrontal cortex (mPFC)-amygdala functional connectivity (Nashiro et al., 2022; Sakaki et al., 2016). Furthermore, compared to older adults in this sample, young adults with higher HRV were discovered to have stronger amygdala-ventrolateral prefrontal cortex (vlPFC) connectivity (Sakaki et al., 2016). Relatedly, a study conducted by Kumral et al. (2019) found that young adults with higher resting HRV exhibited stronger bilateral ventromedial prefrontal cortex (vmPFC) connectivity, with this vmPFC seed demonstrating further extended functional connectivity with several CAN regions. Increasing HRV via biofeedback (e.g., slow breathing, Lehrer & Gevirtz, 2014) has also been shown to elevate resting-state functional connectivity of the prefrontal cortex to neural regions implicated in emotional processing (Schumann et al., 2021a). Specifically, increasing HRV via an 8-week HRV biofeedback intervention was reported to enhance resting-state functional connectivity between the vmPFC and various regions outlined in the NIM, including the amygdala, middle cingulate cortex, anterior insula, and lateral PFC (Schumann et al., 2021a). Interestingly, only a few studies to date have assessed HRV and associated neural activity during tasks that require emotional or self-regulatory processes. One study discovered that higher resting HRV was related to increased vmPFC activation during an effortful self-control dietary task in young adults (Maier & Hare, 2017). Using a voluntary emotion regulation paradigm, Steinfurth et al. (2018) reported that young adults with higher HRV more effectively recruited the dorsal medial prefrontal cortex to modulate amygdala responses via reappraisal. In summary, many of the brain areas identified in HRV neuroimaging studies overlap with regions involved in supporting automatic and voluntary emotion regulatory processes (Morawetz et al., 2020; Wager et al., 2008).

Nonetheless, it is evident that previous research has largely focused on HRV and neural functional connectivity predominantly during rest (i.e., in the absence of an explicit task), with considerably fewer studies focusing on explicit emotion regulation. Resting-state paradigms have recently received criticism in the literature, especially in relation to the utility, interpretability and reliability of neural findings observed under resting-state contexts (Finn, 2021). Indeed, the state of ‘rest’ is increasingly being recognised as a ‘task’ in and of itself, with many unconstrained, internal state factors contributing to diverse cognitive states (Finn, 2021). Recent evidence has highlighted the potential advantage of demands imposed by task engagement, and how such demands may constrain underlying neural functional connectivity to reduce variance related to aforementioned internal state factors, in turn increasing sensitivity to detect individual differences of interest (Finn & Bandettini, 2021). Crucially, since the NIM emphasises the role of the inhibitory cortical-subcortical circuitry in supporting adaptive self-regulation, examining HRV and associated functional connectivity in contexts that require active engagement of emotion regulatory processes may help to further our understanding of heart-brain function in supporting emotion regulation.

In the current study, we sought to extend on previous resting-state functional connectivity findings, examining associations between pulse-derived HRV and neural functional connectivity whilst participants actively engaged in a voluntary emotion regulation task in the scanner. On the basis of prior findings, we hypothesised that HRV would be positively associated with functional connectivity between the amygdala and a region of the mPFC previously associated with HRV (Sakaki et al., 2016). Specifically, we predicted that old and young adults with higher HRV would exhibit stronger positive amygdala-mPFC functional connectivity during a cognitive reappraisal task. Given that pulse recordings were obtained concurrently in the scanning session with the reappraisal task, our primary focus was to examine the relationship between HRV and amygdala connectivity in an emotion regulation context, adopting a functional connectivity analysis similar to that performed on resting-state data (e.g., calculating functional connectivity during the reappraisal task). However, for conceptual replication and comparative purposes, we further assessed pulse-derived HRV and associated resting-state functional connectivity acquired during an initial scanning session that took place 1-2 weeks prior to the session where the HRV measures were obtained (further details and results are presented in the Supplementary Material).

## 2. Materials and Method

### 2.1. Participants

Participants in the current study were derived from a larger sample of 91 subjects (71 old adults, 20 young adults) previously recruited as part of an ageing research project (Lloyd et al., 2021; https://openneuro.org/datasets/ds002620). All participants were recruited via the University of Reading’s Older Adult Research Panel and through local poster and newspaper advertisements in Reading. Participants received financial compensation (£7.50 per hour) for their participation. From the overall sample, 74 participants (55 old adults, 19 young adults) had both emotion regulation task-based functional magnetic resonance imaging (fMRI) and pulse data. Figure 1 illustrates the participant selection and exclusion process. Following exclusion, 70 participants (52 old and 18 young adults, aged 18-84 years, *M* age = 58.27 years, SD = 20.33; 51% male) were considered for analyses (see Table 1 for details per age group).

**Figure 1.**
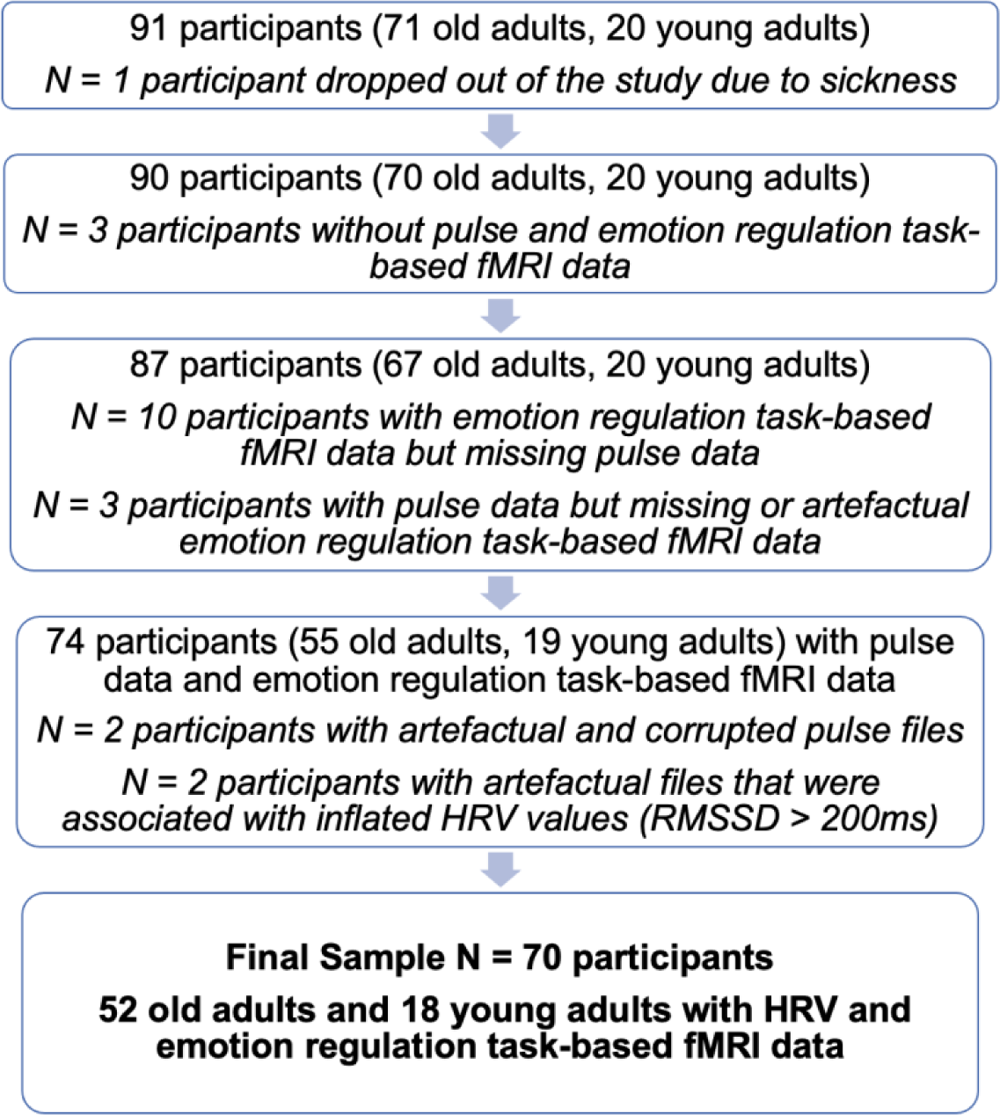
Participant selection and exclusion process. Participants were selected from a larger pool of subjects recruited as part of a wider ageing study.

**Table 1.** Descriptive Statistics for Age, Sex, HRV-Related Metrics and Amygdala-mPFC Connectivity Across the Whole Sample and for Old and Young Adult Sub-Samples

|  | Whole Sample (N = 70)<br>M (SD) | Old Adults (N = 52)<br>M (SD) | Young Adults (N = 18)<br>M (SD) |
| --- | --- | --- | --- |
| Age | 58.27 (20.33) | 69.34 (8.08) | 26.28 (4.75) |
| Age Range | 18 – 84 years | 55 – 84 years | 18 – 35 years |
| Sex (%) | 49% female, 51% male | 44% female, 56% male | 61% female, 39% male |
| InRMSSD (ms) | 4.01 (0.54) | 3.92 (0.55) | 4.29 (0.44) |
| Heart Rate (BPM) | 67.60 (17.64) | 64.61 (9.78) | 76.24 (29.48) |
| RR Interval (ms) | 937.73 (156.19) | 959.80 (141.43) | 873.97 (182.24) |
| Right amygdala-mPFC<br>Connectivity (Parameter estimate) | 0.03 (0.13) | 0.02 (0.11) | 0.05 (0.16) |
| Left amygdala-mPFC Connectivity<br>(Parameter estimate) | 0.03 (0.11) | 0.02 (0.10) | 0.05 (0.11) |

All participants were right-handed and reported no history of neurological disorder. Medical history and medication details were obtained for the older adults only. Of the older adults included in the study (N = 52), 15 disclosed taking regular medication for blood pressure and/or cardiovascular health: statins (N = 8), angiotensin-converting enzyme inhibitors (N = 2), angiotensin receptor blockers (N = 2), calcium channel blockers (N = 2) and beta-blockers (N = 1). The remaining 37 participants did not report use of medication related to cardiovascular health. Furthermore, 21 participants reported having experienced a cardiovascular health condition: high blood pressure (N = 12), high cholesterol (N = 6) and mini-stroke (N = 3). Given that we did not observe significant differences in HRV between those taking cardiovascular medication (*t*(50) = −0.46, *p* = .647, *d* = −0.14) and those who disclosed a history of cardiovascular disease (*t*(50 = −0.70, *p* = .485, *d* = −.20), with participants that did not report use of cardiovascular medication and/or a history of cardiovascular disease, we opted to retain these older adults in the analyses.

The research study from which the current sample was derived was carried out in accordance with the Declaration of Helsinki (1991, p.1194). The study’s procedures were given a favourable ethical opinion of conduct by the University of Reading’s Ethics Committee and NHS Research Ethics Service. Participants provided written informed consent prior to their participation.

### 2.2. Materials and Procedure

#### 2.2.1. Cognitive Reappraisal Task

Participants engaged in a voluntary emotion regulation task during the scan, which followed an established cognitive reappraisal paradigm employed by previous research (e.g., van Reekum et al., 2007). A detailed description of the reappraisal task and stimuli can be found in Lloyd et al. (2021).

The cognitive reappraisal task comprised of 96 trials in total, in which 72 negative and 24 neutral pictures obtained from the International Affective Picture System (IAPS; Lang et al., 2008) were presented. On a given trial, participants were instructed to either “suppress” (decrease), “enhance” (increase), or “maintain” their emotional response and attend to the negative image presented (neutral images were always paired with the “maintain” instruction). The “suppress” instruction involved imagining an outcome less negative than the participant’s original thoughts and/or feelings towards the image, whilst “enhance” required imagining a worse or more negative outcome than originally experienced. In the “maintain” condition, participants were instructed to simply attend to the image and sustain their emotional response. Following the presentation of the picture and engagement in the relevant auditory regulation instruction, participants were asked to rate the picture via a 4-button MR-compatible button box held in the participant’s right hand.

The scanning procedure was distributed across four identical runs, with 24 trials in each run. The duration of each run was approximately 7 minutes, with rest breaks offered between runs, leading to an overall task duration of approximately 30 minutes.

#### 2.2.2. General Procedure

Participants were invited to attend two different sessions within the Centre for Integrative Neuroscience and Neurodynamics (CINN) at the University of Reading. The first session comprised an initial scanning protocol to obtain anatomical T1-weighted (T1w) structural scans, localisers and a resting-state scan, whereby participants were instructed to maintain their gaze on a fixation cross presented in the middle of the screen (rsfMRI scan duration of 10 minutes, 11 seconds). Participants also engaged in several cognitive tasks outside of the scanner which are summarised elsewhere (Lloyd et al., 2021). The first session had an overall duration of approximately three hours (1 hour scanning time). Participants were invited back for a second session which took place a few days (two weeks maximum) after the first session. In the second session, further anatomical (T1w) scans were acquired and participants performed two tasks whilst in the scanner: the cognitive reappraisal (emotion regulation) task and an emotional faces processing task. The participant’s pulse was recorded throughout the scan. The overall duration of the second session was approximately two hours (1 hour scanning time).

### 2.3. Data Reduction and Analysis

#### 2.3.1. HRV Processing and Analysis

A pulse signal was continuously recorded via an MRI-compatible pulse oximeter clip attached to the participant’s left finger throughout the scanning session, including breaks (sampling rate = 50 Hz). The pulse oximeter was integrated with the Siemens Magnetom Trio MRI scanner, from which the raw pulse signal was subsequently extracted.

The raw pulse files underwent visual inspection for quality and usability prior to pre-processing and were formatted to read into LabChart software (version 8.1.11; AD Instruments, Oxford, UK). Initial manual edits within LabChart involved cutting the beginning and/or end of the file where flatlines and/or obvious calibration and motion-related noise were visually detected. Subsequently, LabChart files were converted and exported into LabChart text files to ensure compatibility with Kubios HRV Analysis software (version 2.2; Biosignal Analysis and Medical Imaging Group, University of Kuopio, Finland; Tarvainen et al., 2014). Further processing of the pulse signal and calculation of HRV measures were performed within Kubios. Taking into consideration variation in breaks between runs and tasks, alongside the quality of the pulse signal, participants had somewhat varying durations of pulse signal for analysis (range 17-76 minutes, *M* duration = 51 minutes). Occasionally, the automated peak detection feature either misplaced or missed the peak, thus resulting in manual corrections to either place or (re)move markers to the peak of the pulse waveform. Following manual corrections, data were artefact-corrected using the “low” threshold setting (350 ms) across all participants to retain as many natural variations between heart beats as possible.

The Root Mean Square of Successive Differences (RMSSD), measured in milliseconds, and High-Frequency HRV (HF-HRV), defined using a frequency band of 0.15 – 0.40 Hz, measured in absolute power (ms^2^, Fast Fourier Transform) were calculated within Kubios. Both measures were natural log transformed (ln) to correct for positive skew within RStudio (version 1.4.1106) using the ‘log’ command from the *base* package (v3.5.2). Despite variation in pulse duration, this did not demonstrate a significant correlation with either raw RMSSD (*r* = −.04, *p* = .747) or natural log transformed RMSSD (*r* = .02, *p* = .840) values across participants (N = 70). Whilst RMSSD and HF-HRV metrics reflect parasympathetic vagal control, the RMSSD is a primary and robust measure of vagal tone (Kleiger et al., 2005), that is generally less susceptible to physiological noise, including respiratory influence (Hill et al., 2009). Also, given that both natural log transformed HRV measures exhibited a strong positive association in the current study (*r* = .98, *p* < .001), we proceeded with the (ln)RMSSD as our primary HRV metric for all analyses.

#### 2.3.2. MRI Image Acquisition

Structural and blood oxygenation level dependent (BOLD) functional imaging data were acquired using a 3T Siemens Magnetom Trio MRI scanner with a 12-channel head coil (Siemens, Healthcare, Erlangen, Germany) contained within the CINN at the University of Reading. For each participant, a 3D structural MRI was obtained via a T1-weighted sequence (Magnetization Prepared Rapid Acquisition Gradient Echo (MPRAGE)), repetition time (TR) = 2020 ms, echo time (TE) = 3.02 ms, inversion time (TI) = 900 ms, flip angle 9°, field of view (FOV) = 250 x 250 x 192 mm, resolution = 1 mm isotropic, acceleration factor = 2, averages = 2, acquisition time = 9 minutes, 7 seconds). The emotion regulation fMRI data were obtained in four blocks of identical procedure, using an echo planar imaging (EPI) sequence (211 whole-brain volumes, 30 sagittal slices with P>A phase encoding, slice thickness = 3.0 mm, slice gap = 33%, TR = 2000 ms, TE = 30 ms, flip angle = 90°, FOV = 192 x 192 mm^2^, resolution = 3 mm isotropic, acquisition time = 7 minutes 10 seconds per block). The structural and emotion regulation fMRI task data are publicly available on OpenNeuro: https://openneuro.org/datasets/ds002620/versions/1.0.0.

#### 2.3.3. MRI Data Pre-processing

Functional imaging data were pre-processed and analysed using FMRIB’s Software Library (FSL, version 6.0; www.fmrib.ox.ac.uk/fsl; Jenkinson et al., 2012; Woolrich et al., 2009; Smith et al., 2004) and Analysis of Functional NeuroImages (AFNI, version 19.3.03; http://afni.nimh.nih.gov/afni; Cox, 1996). Initial pre-processing steps included: skull stripping (non-brain removal) using FSL’s brain extraction tool (BET; Smith, 2002), motion correction using MCFLIRT (Jenkinson et al., 2002), field-map correction to correct for potential magnetic field inhomogeneity distortions, spatial smoothing using a Gaussian kernel with a full-width half maximum (FWHM) of 5 mm and high-pass temporal filtering (Gaussian-weighted least squares straight line fitting with sigma = 50 s). Each subject’s native image was normalised to Montreal Neurological Institute (MNI) space via co-registration to their high resolution T1-weighted image.

Application of FSL’s MELODIC Independent Components Analysis (ICA; Beckmann & Smith, 2004) separated the fMRI BOLD signal into a set of spatial maps (independent components) representing neural signal and/or noise. Independent components containing structured temporal noise, including scanner and hardware artefacts, physiological artefacts (respiratory and/or cardiac noise), and motion-related noise were identified via visual inspection and removed using the FSL command line tool ‘*fslregfilt*’ for each emotion regulation task run (Griffanti et al., 2017). An average percentage of 72.07% components were removed across the four runs. This is generally in line with previous research that has typically identified >70% noise versus signal components in standard sequences at 3T (Griffanti et al., 2017).

Following ICA filtering, low bandpass filtering was applied to the fMRI data using AFNI’s ‘*3dBandpass*’ tool (Cox, 1996) to further remove confounding signals below 0.009 Hz and above 0.1 Hz. Prior to analysis, each subject’s corresponding mean functional timeseries image was added back to the bandpass filtered data using *fslmaths* to ensure compatibility with FSL’s FMRI Expert Analysis Tool (FEAT).

#### 2.3.4. Functional Connectivity Analysis

Regions of interest (ROIs) were separate right and left amygdala seeds, and an area of the mPFC previously found to be correlated with HRV (Sakaki et al., 2016). Separate amygdala ROIs were selected given recent discrepancies in amygdala lateralisation with the mPFC as a function of HRV (Nashiro et al., 2022; Sakaki et al., 2016), and also observed lateralisation effects highlighted in previous research concerning emotion processing and regulation (Baas et al., 2004; Yang et al., 2020). Amygdala ROI masks were defined using the Harvard-Oxford Subcortical Probability atlas and thresholded at 80% probability. The mPFC ROI employed by Sakaki et al. (2016) and in the present study was derived from a significant cluster previously correlated with memory positivity (Sakaki et al., 2013), containing voxels from anterior cingulate cortex (ACC) and paracingulate gyrus (Harvard-Oxford atlas), thresholded at 25% probability.

All ROI masks (right and left amygdala, mPFC) were first transformed to each participant’s native functional space using FSL’s Apply FLIRT Transform ‘*ApplyXFM*’ and binarised. Subsequently, the mean time series for each ROI was extracted from the four separate emotion regulation runs for each participant using ‘*fslmeants*’.

Separate first-level regression analyses were performed for each ROI using FEAT (Woolrich et al., 2001). Similar to a functional connectivity analysis typically performed on resting-state data, individual FEAT models included the mean time series extracted from the specific ROI and regressors of no interest, specifically: FSL’s six standard head-motion parameters, and average white matter and ventricular (CSF) signal. Average signal from white matter and CSF was extracted from masks generated via segmentation of each participant’s high resolution T1w image using FSL’s FAST algorithm (Zhang et al., 2001).

Inclusion of global signal regression (GSR) has received scrutiny in the literature (Murphy et al., 2009; Murphy & Fox, 2017; Uddin, 2017). Given the controversy and lack of consensus surrounding GSR, we decided not to include GSR as a regressor in the model. Importantly, we did not include the task design as a regressor in our model either. It is recognised that not including the task design as a regressor in task-based functional connectivity analyses can result in spurious correlations and systematic inflation of functional connectivity estimates due to task-induced coactivations (Cole et al., 2019). However, the overarching aim of the present study is to examine HRV and associated neural functional connectivity in a voluntary emotion regulation context. Since HRV is closely related to, and considered a metric of, regulatory processes, including the task design as a regressor would remove variance of interest and relevance to the aim of our study. Furthermore, not regressing the task design has been reported to increase the reliability of functional connectivity measures (Cho et al., 2021), whilst other studies have found that inclusion versus non-inclusion of the task design in task-based connectivity analyses does not appear to significantly change the overall pattern of the functional connectivity findings reported (Cao et al., 2018; Finn, 2021; Kraus et al., 2021).

Prior to group-level analyses, a second-level fixed effects analysis using FSL’s FEAT was applied to the emotion regulation task-based fMRI data to collapse the ROI connectivity maps across the four task runs^1^. This generated positive and negative mean contrast of parameter estimates (COPE) images for input to higher-level analyses.

#### 2.3.5. Amygdala-mPFC Functional Connectivity Analyses

Beta values from right and left amygdala (positive COPE) connectivity maps were extracted using FSL’s Featquery, with the mPFC seed as the reference mask. The corresponding mean parameter estimates served as an index of amygdala-mPFC connectivity strength.

#### 2.3.6. Whole-Brain Functional Connectivity Analyses

Given the heterogeneous neurological profiles often observed in ageing brains (Chen et al., 2016), and the larger sample of older adults recruited in the current study, we performed whole-brain functional connectivity analyses for all ROIs across the whole sample, including age as a blocking factor in the analyses, and further performed separate whole-brain analyses restricted to the older adult sample only. This allowed us to be more inclusive in our search for functionally-relevant regions associated with HRV that may have been excluded or otherwise missed using a ROI approach. Furthermore, the decision to run separate whole-brain connectivity analyses restricted to the old adult sample was primarily driven by the unequal number of old relative to young adults (and the comparative small sample size of the young adult group), along with the strong effect of biological age on HRV (Agelink et al., 2001; Russoniello et al., 2013).

Whole-brain group analyses were performed using FSL’s FEAT (Woolrich et al., 2004). Separate FMRIB’s Local Analysis of Mixed Effects (FLAME) whole-brain analyses were carried out for each seed region. The general linear model (GLM) included four explanatory variables: group mean and three predictors, HRV (lnRMSSD, centred), age (effect coded using +1 and −1 to define old and young adult groups respectively) and a HRV by age interaction term (lnRMSSD centred x age group). Seven contrasts were entered into the model: group mean, HRV, age and the HRV by age interaction term (positive and negative contrasts for each EV). Clusters surviving a threshold of Z > 3.1 and correction for multiple comparisons with Gaussian random field theory (cluster significance: *p* = 0.05-corrected) were identified (Worsley, 2001). The locations of significant clusters that survived correction were labelled using the Harvard Oxford Cortical Structural and Subcortical atlases in MNI space within FSL. Mean parameter estimate (beta) values from significant clusters that emerged as a main effect of HRV were extracted for visualisation purposes.

## 3. Results

### 3.1. Descriptive Statistics

Table 1 summarises general descriptives for the whole sample and for old and young adult age groups separately. HRV significantly differed by age group, such that older adults demonstrated significantly reduced HRV as indexed by lower (ln)RMSSD values (*M* = 3.92, *SD* = 0.55), in comparison to young adults (*M* = 4.29, *SD* = 0.44), *F*(1,66) = 6.06, *p* = .016, η_p_^2^ = 0.08. However, there was no significant difference in (ln)RMSSD values between females (*M* = 4.07, *SD* = 0.52) and males (*M* = 3.96, *SD* = 0.57) across the whole sample, *F*(1,66) = 0.09, *p* = .764, η_p_^2^ = 0.00, nor was there a significant interaction between age group and sex on (ln)RMSSD values, *F*(1,66) = 0.15, *p* = .698, η_p_^2^ = 0.00. Thus, no significant differences in HRV related to sex were observed in the present study (see Figure S1 in the Supplementary Material). Additionally, there was a significant difference in the mean RR interval (*t*(68) = 2.06, *p* = .044, *d* = 0.56), but no significant difference in mean heart rate (*t*(18) = −1.64, *p* = .117, *d* = −0.68) between old and young adults.

### 3.2. HRV and Amygdala-mPFC Functional Connectivity Analysis

Multiple regression analyses were employed to examine associations between HRV and amygdala-mPFC functional connectivity strength in the whole sample. Separate multiple regression models were tested with (i) right amygdala-mPFC connectivity and (ii) left amygdala-mPFC connectivity values as dependent variables. A segregation in age (years) was observed between the old and young adults, leading to a natural formation of two separate age groups (see Figure S2 in the Supplementary Material). We therefore entered age as a categorical predictor in the regression models. The following predictors were entered into the regression model: age group (1 = old adults, 0 = young adults), (ln)RMSSD (centered), and a HRV x age interaction term^2^. In each model, age group and HRV were entered first (step 1), followed by the HRV x age interaction predictor (step 2). Standardised beta coefficients are reported for all predictors.

#### 3.2.1. HRV and Right Amygdala-mPFC Functional Connectivity

Neither age (*β* = −0.12, *t* = −0.94, *p* = .350) or HRV (*β* = −0.02, *t* = 0.14, *p* = .886) contributed significantly to the overall regression model, *F*(2,67) = 0.45, *p* = .637, explaining only 1.3% of the variance in right amygdala-mPFC functional connectivity. Entering the HRV x age interaction term into the model improved the proportion of variance explained in right amygdala-mPFC connectivity (Δ*R*^2^ = 0.13, *F*(3,66)= 3.62, *p* = .018). The interaction between HRV and age was found to significantly predict right amygdala-mPFC functional connectivity strength (*β* = 0.86, *t* = 3.14, *p* = .003). Follow-up regression models indicated that the younger adults appeared to drive this interaction, such that young adults with higher HRV exhibited significantly weaker right amygdala-mPFC functional connectivity (*β* = −0.54, *t* = −2.54, *p* = .022), whereas old adults demonstrated a slight positive, albeit non-significant, association between HRV and right amygdala-mPFC connectivity during the task (*β* = 0.17, *t* = 1.24, *p* = .222) (Figure 2a).

**Figure 2.**
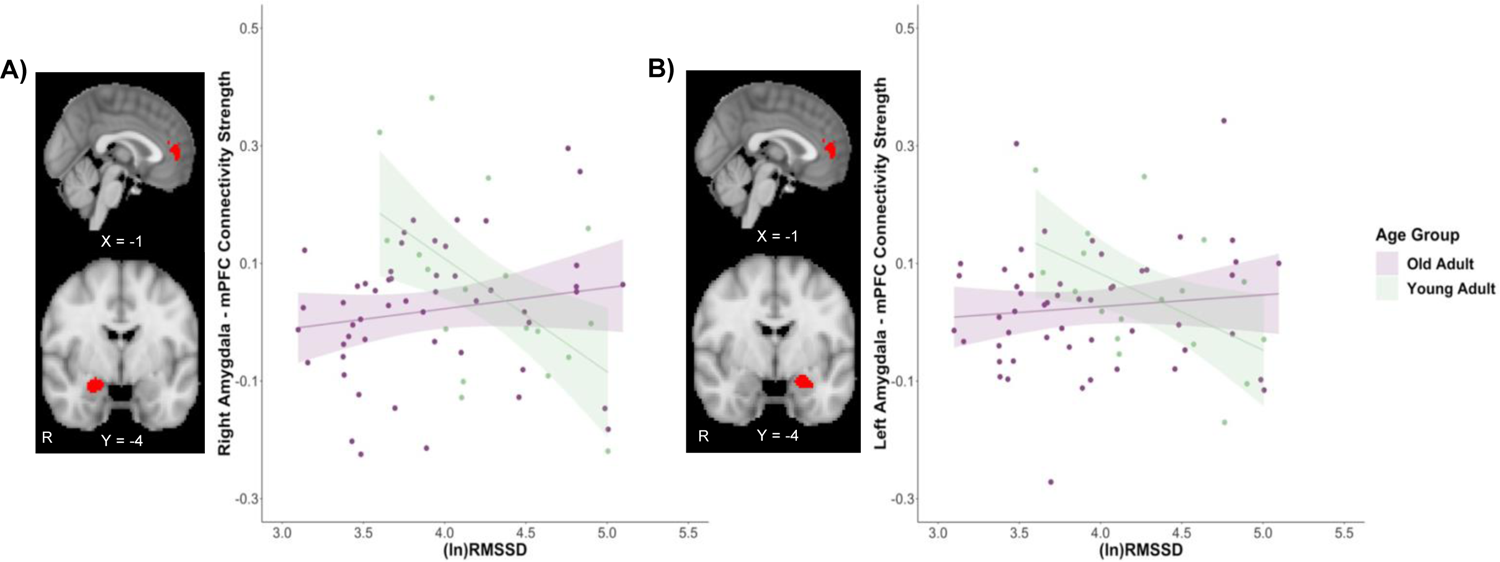
HRV and amygdala-mPFC functional connectivity during the reappraisal task. **A)** mPFC seed (top) and right amygdala seed (bottom). Significant HRV x age interaction for right amygdala-mPFC connectivity strength. In young adults (light green), higher HRV significantly predicted weaker connectivity between the right amygdala and mPFC, whereas a slight positive, albeit non-significant, association between HRV and right amygdala-mPFC connectivity was observed in the old adults (purple). **B)** mPFC seed (top) and left amygdala seed (bottom). Significant HRV x age interaction for left amygdala-mPFC connectivity strength. Similar to the right amygdala connectivity findings, in young adults, greater HRV significantly predicted weaker left amygdala- mPFC connectivity, whereas a non-significant, weak positive association between HRV and left amygdala- mPFC connectivity was observed in the old adults during the reappraisal task. *(ln)RMSSD*; natural log transformed root mean square of successive differences.

#### 3.2.2. HRV and Left Amygdala-mPFC Functional Connectivity

Similar to the right amygdala-mPFC functional connectivity findings, HRV (*β* = −0.03, *t* = −0.26, *p* = .797) and age (*β* = −0.09, *t* = −0.73, *p* = .466) did not contribute significantly to the overall model, *F*(2,67) = 0.27, *p* = .764, and explained very minimal variance (0.8%) in left amygdala-mPFC functional connectivity strength. However, when the HRV x age interaction term was entered into the model, this improved the proportion of variance explained in left amygdala-mPFC connectivity (Δ*R*^2^ = 0.08), although the overall model remained non-significant, *F*(3,66) = 2.07, *p* = .112. The HRV x age interaction was found to predict left amygdala-mPFC connectivity strength (*β* = 0.67, *t* = 2.38, *p* = .020). Follow-up regression models per age group revealed younger adults to drive this significant interaction, whereby greater HRV significantly predicted weaker left amygdala-mPFC functional connectivity in young adults (*β* = 0.51, *t* = −2.37, *p* = .031). Conversely, a non-significant, weak positive association between HRV and left amygdala-mPFC connectivity strength was observed in old adults (*β* = 0.10, *t* = 0.74, *p* = .461) (Figure 2b).

### 3.3. Whole-Brain Functional Connectivity Analyses

#### 3.3.1. Right Amygdala Whole-Brain Functional Connectivity

Significant clusters surviving correction as a main effect of HRV for the right amygdala whole-brain functional connectivity analyses are displayed in Table 2. Across old and young adults, higher HRV was associated with weaker right amygdala connectivity between the right angular gyrus (extending into right superior lateral occipital cortex), and bilateral posterior cingulate gyrus (Z > 3.1, *p* = 0.05-corrected). A scatterplot displaying beta values extracted from the bilateral posterior cingulate gyrus cluster with HRV are displayed in Figure 3. No other clusters survived correction for the positive HRV contrast, nor for positive or negative HRV by age interaction contrasts across the whole sample.

**Table 2.**
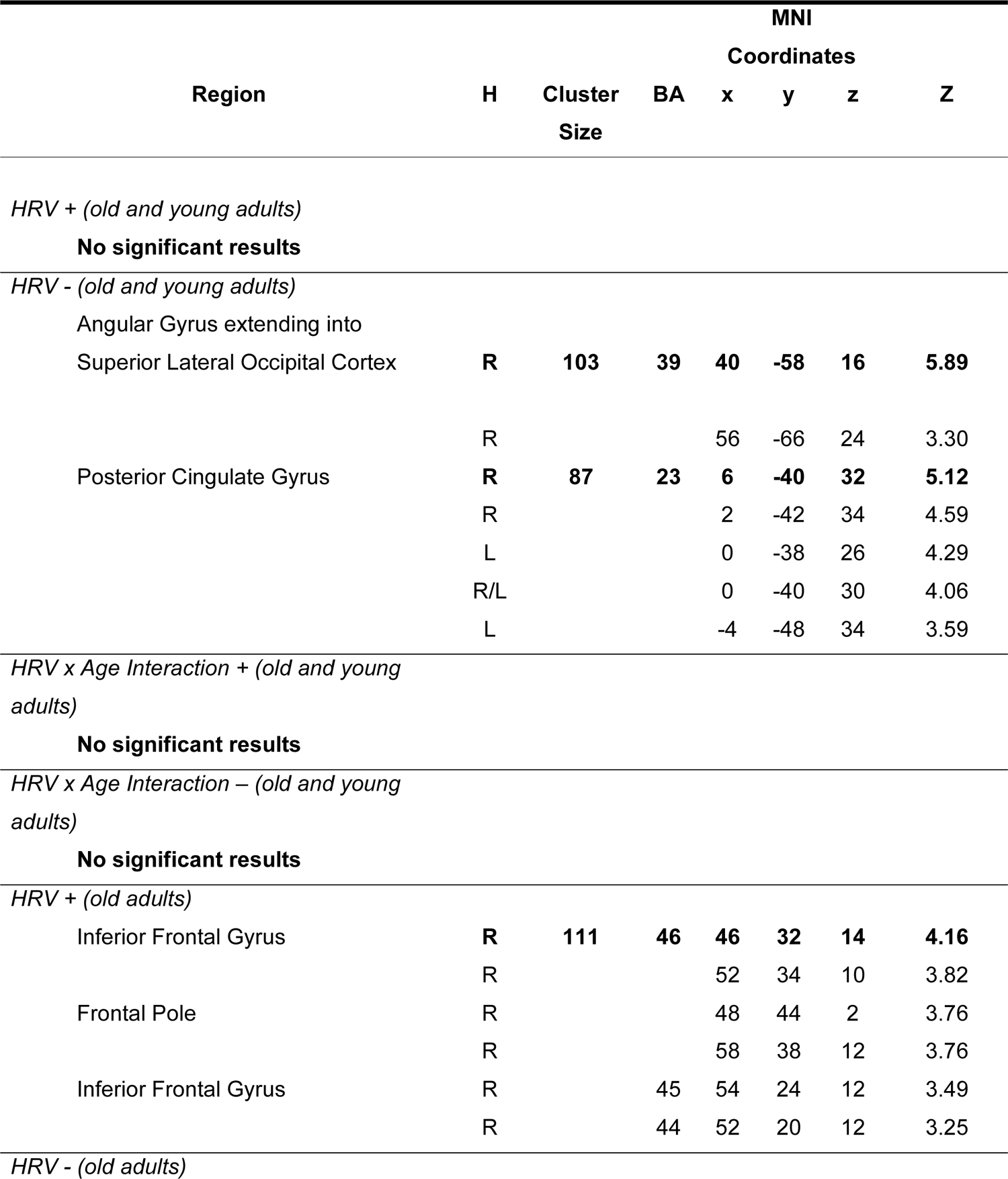
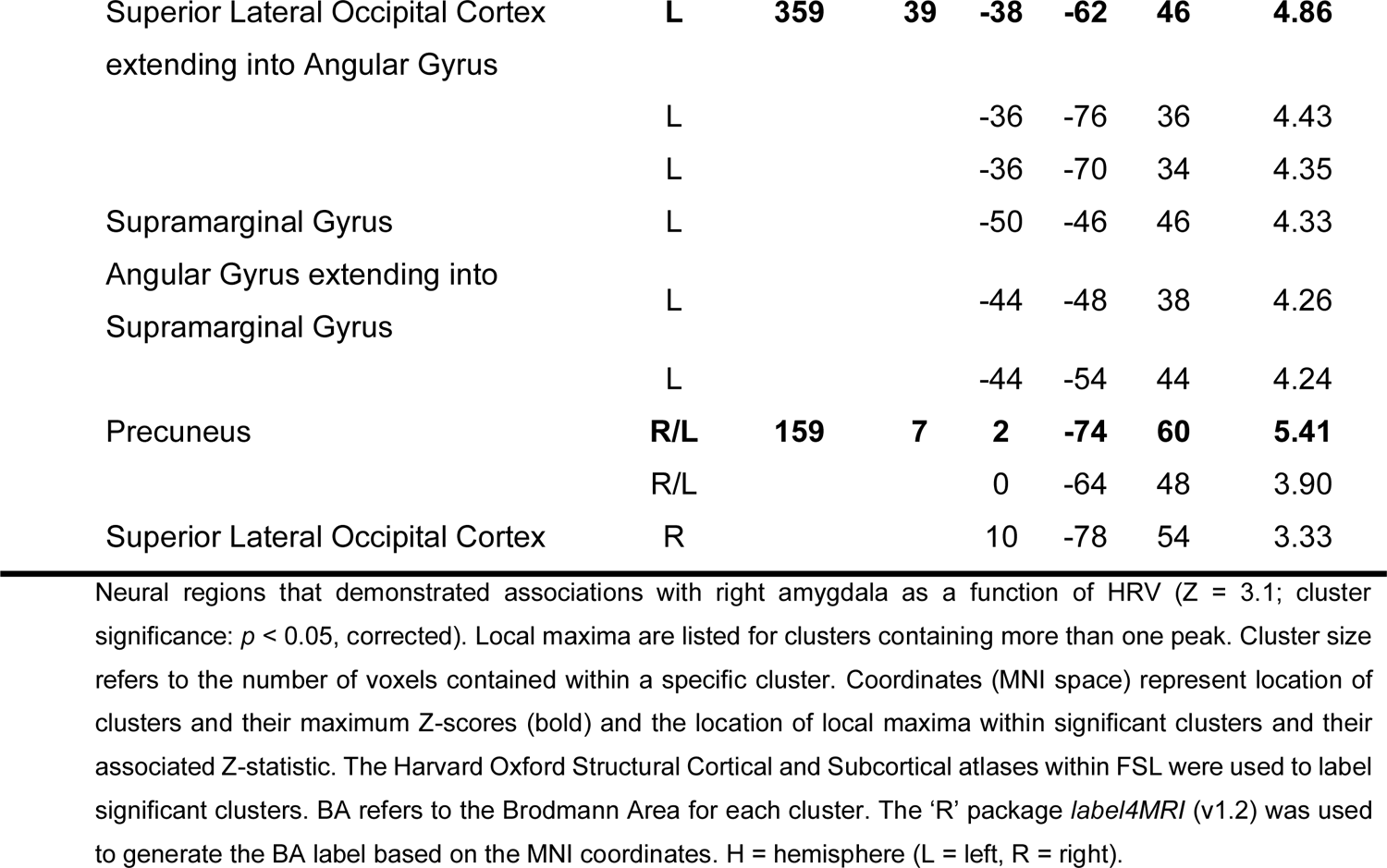
Neural Regions and Local Maxima for Right Amygdala Whole-Brain Connectivity

**Figure 3.**
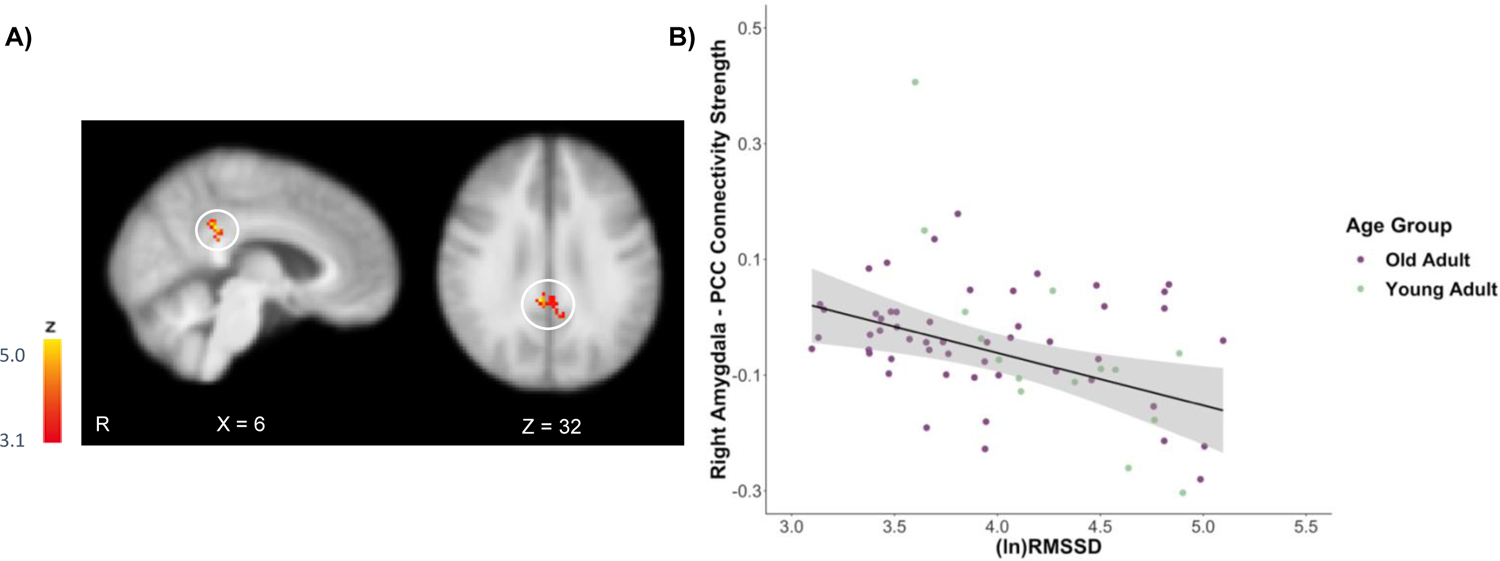
**A)** Significant bilateral posterior cingulate cortex (PCC) cluster that survived correction as a main effect for the negative HRV contrast in the right amygdala whole-brain analysis (Z > 3.1, *p* = 0.05-corrected). **B)** Scatterplot displays the inverse association between HRV ((ln)RMSSD) values and standardised beta values depicting right amygdala- bilateral PCC connectivity strength in the whole sample during the reappraisal task in old and young adults (N = 70). Note that the different colours assigned to old (purple) versus young (light green) adult age groups are depicted for display purposes only. *(ln)RMSSD*; natural log transformed root mean square of successive differences.

Repeating this analysis on the old adult sample only, a significant main effect of HRV emerged, such that higher HRV was positively correlated with stronger functional connectivity between the right amygdala and the right inferior frontal gyrus, a cluster forming part of the right ventrolateral prefrontal cortex (vlPFC). A scatterplot displaying beta values extracted from this right vlPFC cluster with HRV are displayed in Figure 4. Moreover, for the HRV negative contrast, higher HRV was associated with weaker right amygdala connectivity with several regions, including bilateral superior lateral occipital cortex extending into left angular and supramarginal gyrus, and bilateral precuneus.

**Figure 4.**
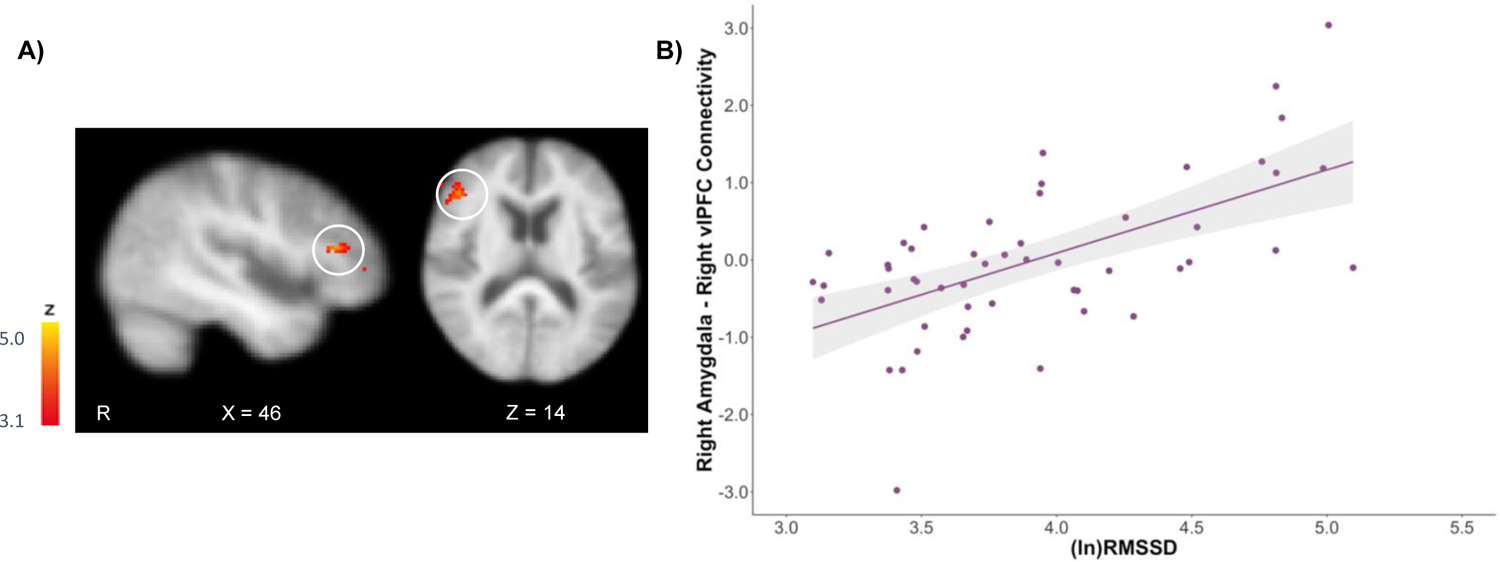
**A)** Significant right inferior frontal gyrus (vlPFC) cluster that survived correction as a main effect for the positive HRV contrast in the right amygdala whole-brain analysis restricted to the old adult sample (Z > 3.1, *p* = 0.05-corrected). **B)** Scatterplot displays the positive association between HRV ((ln)RMSSD) values and standardised beta values depicting right amygdala - right vlPFC connectivity strength in the old adult sample (controlling for age). *(ln)RMSSD*; natural log transformed root mean square of successive differences.

#### 3.3.2. Left Amygdala Whole-Brain Functional Connectivity

No significant clusters survived correction as a function of HRV for left amygdala functional connectivity in the whole sample (Z > 3.1, *p* = 0.05-corrected), suggesting that HRV did not covary with left amygdala whole-brain functional connectivity across old and young adults throughout the reappraisal task.

When the left amygdala voxelwise whole-brain search was restricted to the old adult sample, a significant positive main effect of HRV was observed, in which higher HRV was correlated with stronger left amygdala connectivity with the right inferior frontal gyrus (vlPFC) and more extensively with the right precentral gyrus (Z > 3.1, *p* = 0.05-corrected). Furthermore, significant clusters also survived correction for the negative HRV contrast, such that higher HRV correlated with reduced left amygdala – left lateral occipital cortex connectivity. Other brain regions that survived correction as a main effect of HRV for the left amygdala whole-brain functional connectivity analyses in the old adults are displayed in Table 3.

**Table 3.**
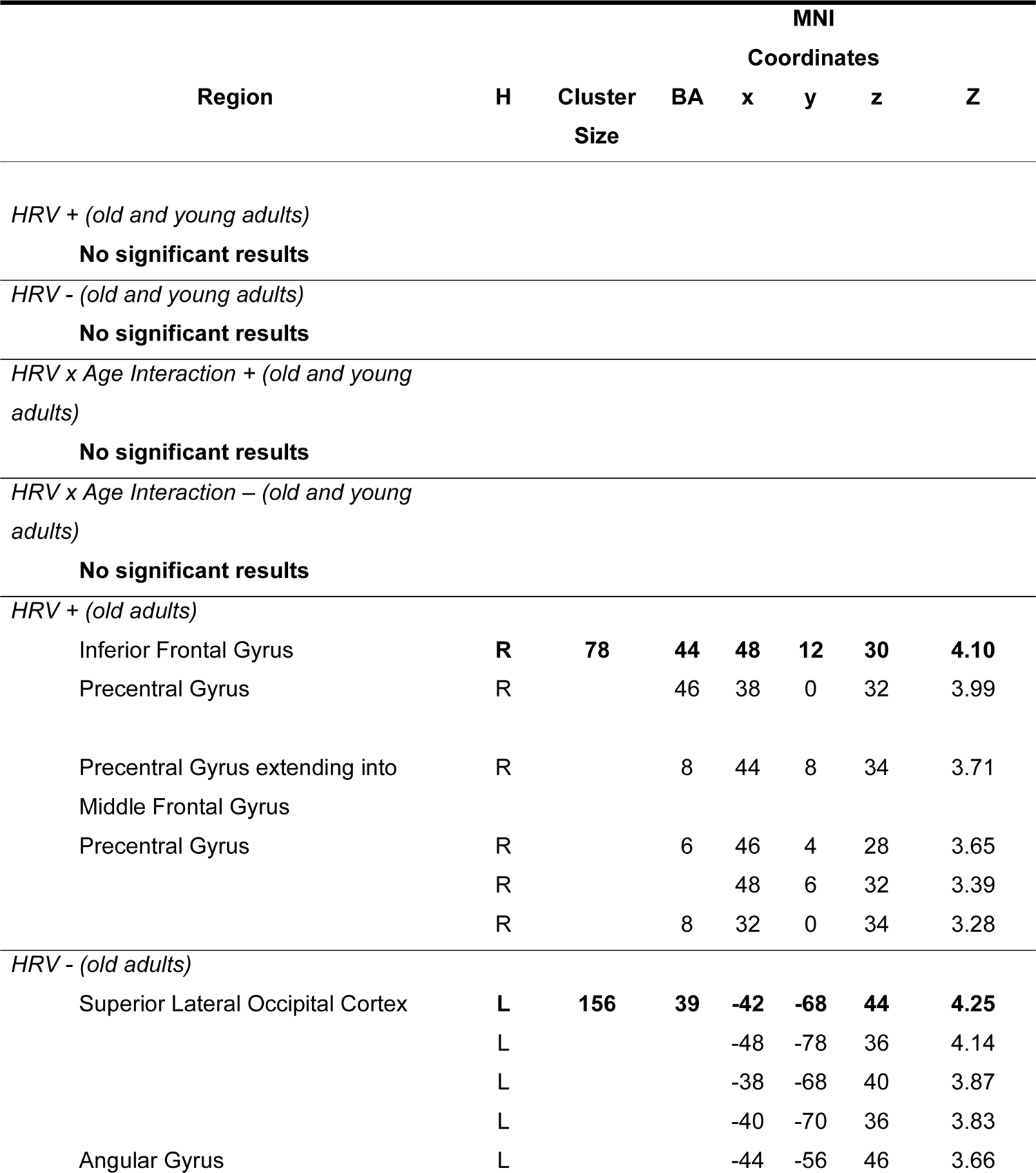
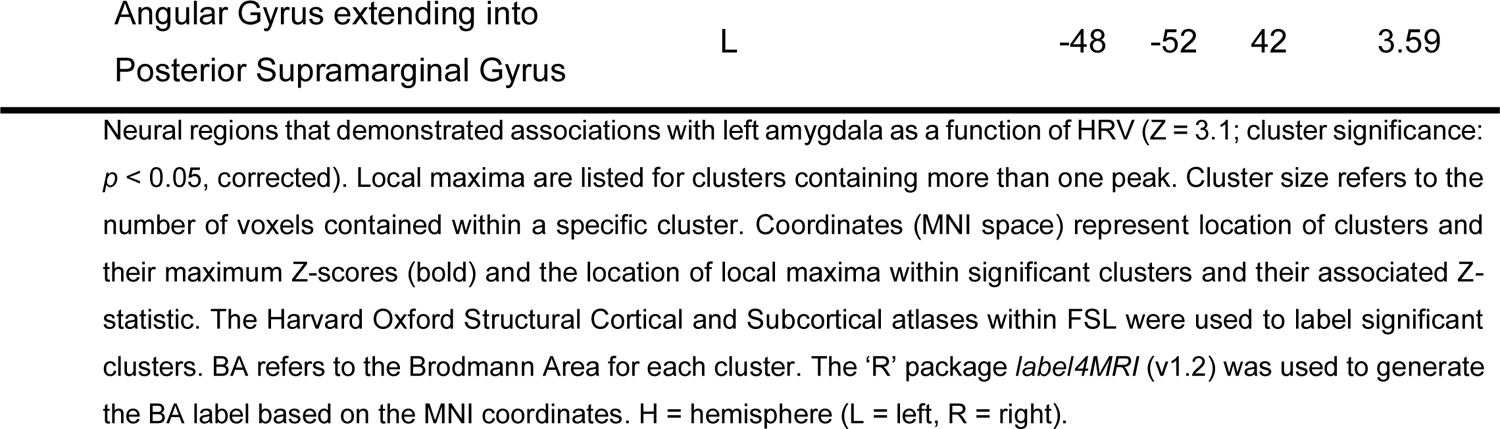
Neural Regions and Local Maxima for Left Amygdala Whole-Brain Connectivity

#### 3.3.3. MPFC Whole-Brain Functional Connectivity

No clusters survived correction as a main effect of HRV for the mPFC seed in a voxelwise whole-brain search in the whole sample, nor when the analysis was restricted to the old adult sample (Z > 3.1, *p* = 0.05-corrected). Therefore, HRV did not significantly predict functional connectivity of this particular area of the mPFC during the reappraisal task.<colcnt=1>

### 3.4. Resting-state Functional Connectivity

In this study, resting-state fMRI was collected approximately a week prior to the emotion regulation task and associated HRV, which is not optimal to infer HRV-resting state associations. Nonetheless, for the purpose of comparing results in the task-based data to resting-state, and to findings reported previously by others (Kumral et al., 2019; Nashiro et al., 2022; Sakaki et al., 2016), we have included the amygdala-mPFC functional connectivity findings and a full report of whole-brain connectivity with the BOLD response in left and right amygdala ROIs in the Supplementary Material (see Tables S1-S3). The whole-brain connectivity results partially replicated prior findings, with higher HRV associated with positive left amygdala-mPFC coupling, albeit that this association did not survive thresholding or correction for multiple comparisons.

## 4. Discussion

The principal aim of the present study was to examine the relationship between HRV and neural functional connectivity whilst old and young adults engaged in a voluntary emotion regulation task. Based on the NIM (Smith et al., 2017; Thayer & Lane, 2000, 2009), we hypothesised that higher HRV would positively correlate with stronger functional coupling between the amygdala and mPFC in an active regulatory context. In old adults, we observed a slight positive, but non-significant, association between HRV and amygdala-mPFC connectivity, partially supporting our hypothesis. Conversely, young adults displayed a stronger, inverse association, whereby higher HRV was linked to reduced functional connectivity between the amygdala and mPFC. Furthermore, in a voxelwise whole-brain search, we discovered that old and young adults with higher HRV exhibited weaker right amygdala-PCC connectivity. Interestingly, in old adults, higher HRV was associated with stronger coupling between the right amygdala and right vlPFC. Our findings indicate that HRV covaries with amygdala functional connectivity during emotion regulation, and more crucially highlight the importance of assessing HRV and brain function during an active emotion regulatory context.

Functional connectivity between the amygdala and mPFC is proposed to support adaptive emotion regulation, with HRV posited to serve as a peripheral index of prefrontal inhibitory control (Thayer & Lane, 2000, 2009; Thayer et al., 2009a). In line with this proposition, prior studies have reported positive associations between HRV and amygdala-mPFC connectivity strength irrespective of age (Nashiro et al., 2022; Sakaki et al., 2016). However, within the context of the emotion regulation task, we found significant interactions between age and HRV to predict both right and left amygdala coupling with the mPFC. The direction of the effect was unexpected, with the young adults driving the interaction, but in whom higher HRV was linked to weaker, rather than a strong positive, coupling between the amygdala and mPFC. It is possible that this particular region of the mPFC is more heavily recruited during rest, compared to an active emotion regulation context. Indeed, during rest, we found a sub-threshold cluster within the mPFC close to our ROI that demonstrated increased functional connectivity with the left amygdala as a function of higher HRV across old and young adults (see Figure S3 in the Supplementary Material). Recently, Nashiro et al. (2022) also found that increases in HRV via biofeedback were correlated with stronger left, but not right, amygdala coupling with the mPFC at rest. Furthermore, prior work has found inverse amygdala-mPFC coupling when using reappraisal to decrease negative affect in a student-aged population (Lee et al., 2012). Hence, the inverse association reported here in young adults may be driven by the decrease conditions throughout the task, although at this stage these findings would require replication using a more targeted event-related connectivity analysis, which is beyond the scope of this manuscript. Taken together, our findings potentially suggest that the regulatory context can affect both the laterality and directionality of amygdala-mPFC functional connectivity associations with HRV.

Furthermore, higher HRV was significantly associated with weaker right amygdala connectivity between the right angular gyrus and bilateral PCC across the emotion regulation task in old and young adults. Both the angular gyrus and PCC form major nodes of the default mode network (DMN), a neural hub implicated in autobiographical memory (Buckner & Carroll, 2007), and self-referential processing (Raichle et al., 2001). Weaker resting-state functional connectivity between the right amygdala and PCC has previously been linked to greater reappraisal success (i.e., effective down-regulation of negative emotion) in young adults (Uchida et al., 2015), whereas increased amygdala-PCC resting-state functional connectivity has been observed following exposure to an acute stressor (Veer et al., 2011). More recently, Baez-Lugo et al. (2021) reported that greater right amygdala-PCC functional connectivity following exposure to videos containing highly negative emotional content (i.e., people suffering) was significantly correlated with higher rumination, anxiety and stress in elderly individuals (Baez-Lugo et al., 2021). Critically, those older adults who self-reported more frequent negative thoughts while watching the negative emotional videos were those who also exhibited stronger right amygdala-PCC connectivity. Considering that lower HRV has been linked to both increased rumination and emotion dysregulation (Visted et al., 2017; Williams et al., 2017), the observation of weaker right amygdala-PCC connectivity in old and young adults with higher HRV in our study may therefore reflect an increased ability to effectively engage with the emotion regulation task at hand.

Finally, we found that older adults with higher HRV exhibited stronger functional connectivity between the amygdala and right vlPFC in a reappraisal context. This finding is particularly interesting since Sakaki et al. (2016) reported a similar association between HRV and amygdala-vlPFC connectivity during rest in young, but not old adults, suggesting that young adults with higher HRV were more likely to spontaneously recruit neural regions involved in explicit emotion regulation. The vlPFC has increasingly been identified as a pivotal neural region involved in emotion regulatory processes (Wager et al., 2008; Zhao et al., 2021), and is an area in which age-related differences have been reported during reappraisal (Opitz et al., 2012; Winecoff et al., 2011). The vlPFC, and lateral prefrontal cortex more broadly, is particularly vulnerable to structural and functional atrophy in healthy ageing (Fjell et al., 2009; Raz et al., 2004). The present finding suggests that higher HRV in older age, at least in a voluntary emotion regulation context, may support increased engagement, and possibly functional preservation, of lateral prefrontal cortex, specifically the right vlPFC, facilitating effective response inhibition and reappraisal of negative emotions. Although the left vlPFC has been more frequently reported in reappraisal studies (Berboth & Morawetz, 2021; Buhle et al., 2014), involvement of the right vlPFC here may be characterised by dominance of the right hemisphere in supporting inhibitory-related processes for affective, cognitive and physiological regulation more broadly (Lane et al., 2009; Thayer et al., 2009b, 2012). Irrespective of any laterality, our findings build on the extant literature on prefrontal mechanisms in reappraisal by highlighting that elevated HRV is associated with positive coupling between the amygdala and vlPFC, which may have implications for psychological wellbeing and resilience in later life.

A few important limitations should be considered when interpreting our findings. Our sample comprised a larger pool of old relative to young adults, leading to an unequal age distribution. Although age was included as a predictor in our regression models, the small sample of young adults rendered any findings specific to the young group as possibly spurious and requiring replication in a larger sample. Furthermore, HRV was derived from a finger pulse oximeter whilst participants were lying down in the scanner and whilst engaging in emotion-related tasks, predominantly reappraisal. Both factors have previously been shown to elevate heart rate and HRV (Butler et al., 2006; Cacioppo et al., 1994), and the use of photoplethysmography to derive HRV metrics, especially RMSSD (Schumann et al., 2021b), could have further resulted in a higher HRV estimate. Additionally, other lifestyle factors known to influence HRV measures, including smoking status, general fitness/activity level, caffeine intake and body mass index (Hayano et al.,1990; Karason et al., 1999; Sammito & Böckelmann, 2016) were not obtained, therefore we cannot rule out the influence of these factors on the current findings. Future research should aim to acquire reliable heart rate recordings to derive HRV metrics both inside and outside of the scanner (Schumann et al., 2021b), alongside potential aggregation of HRV measures across contexts, to capture variance that more strongly represents ‘trait-like’ HRV (see Bertsch et al., 2012).

Whilst our study augments prior findings which have heavily relied on associations between HRV and functional connectivity during rest by assessing heart-brain function in an active emotion regulatory context, the current study and the majority of prior work have typically relied on relatively static functional connectivity techniques. Although a few studies have examined transient HRV changes and functional connectivity using dynamic functional connectivity (dFC) techniques such as the sliding window approach (Chand et al., 2020; Chang et al., 2013; Schumann et al., 2021a), this method is limited by its reliance on arbitrary selection of truncated time windows to assess both functional connectivity and HRV, with the latter particularly affected by the shorter duration of the measurement period (Shaffer & Ginsberg, 2017; TaskForce, 1996). It would therefore be fruitful for future research to employ novel and alternative dFC methods that overcome existing constraints (e.g., co-activation pattern analysis; Liu et al., 2013, 2018) to determine associations between HRV and dynamic neural networks underlying adaptive and flexible regulation across the lifespan.

In conclusion, the current study extends prior resting-state findings by highlighting that HRV covaries with amygdala-cortical functional connectivity in the context of a voluntary emotion regulation task. Particularly, whilst our findings partially replicate amygdala-mPFC connectivity during rest to be coupled to HRV, the task-based covariation between functional connectivity of amygdala-vlPFC and amygdala-PCC and HRV provide further, and more direct, support of the NIM. Furthermore, the findings support the notion that HRV is linked to neural mechanisms that facilitate adaptive emotion regulation, which could have implications for wellbeing and resilience in later life. Collectively, our findings highlight the importance of assessing neurovisceral circuitry during active regulatory contexts to further elucidate core neural mechanisms involved in supporting adaptive self-regulation as a function of HRV more broadly.

## Data Availability Statement

The MRI data that support the findings of this study are openly available on OpenNeuro: https://doi.org/10.18112/openneuro.ds002620.v1.0.0. The pulse data, processing and analysis scripts that support this study are openly available on the Open Science Framework (OSF): https://osf.io/6zdph/.

## Acknowledgements

The authors would like to thank Karis Colyer Patel and Laura Bucher for their assistance with processing the pulse data, Shan Shen for MRI support and help in MRI data acquisition, and all participants for devoting their time to our research. This research was supported by grants from the Biotechnology and Biological Sciences Research Council (BB/J009539/1 and BB/L02697X/1) awarded to Carien van Reekum.

## Footnotes

1 Two participants were missing the final run of the emotion regulation task (run 4), so ROI connectivity maps were averaged across the three available task runs (runs 1-3) for these participants.

2 To reduce the influence of multicollinearity that can occur between the original variables and the subsequent interaction that is comprised of those variables, the HRV x age interaction term was calculated by multiplying the centered (ln)RMSSD scores by the dummy coded age group.

