## Supplementary Material for "Heart Rate Variability Covaries with Amygdala Functional Connectivity During Voluntary Emotion Regulation"

**S1. Resting-state fMRI Details**

*S1.1 Resting-state fMRI Data Acquisition*

Resting-state fMRI data were acquired using an echo planar imaging (EPI) sequence (200 whole-brain volumes, 56 sagittal slices, with P>A phase encoding, slice thickness = 3.0mm, slice gap = 0%, TR = 3000 ms, TE = 30 ms, flip angle = 90°, FOV = 192 x 192 mm^2^, resolution = 3 mm isotropic, acquisition time = 10 minutes, 11 seconds).

*S1.2 Resting-state Scan Procedure*

Participants were asked to maintain their gaze on a fixation cross in the middle of screen presented on a white background. The total duration of the resting-state scan was 10 minutes and 11 seconds.

*S1.3 Resting-state fMRI Pre-processing*

Resting-state functional imaging data were preprocessed and analysed using FMRIB’s Software Library (FSL, version 6.0; Jenkinson et al., 2012; Woolrich et al., 2009; Smith et al., 2004) and Analysis of Functional NeuroImages (AFNI, version 19.3.03; <http://afni.nimh.nih.gov/afni>; Cox, 1996), akin to the preprocessing and analytical procedure applied to the emotion regulation fMRI task-based data. These initial pre-processing steps included: skull stripping (non-brain removal) using FSL’s brain extraction tool (BET; Smith, 2002), motion correction using MCFLIRT (Jenkinson et al., 2002), spatial smoothing using a Gaussian kernel with a full-width half maximum (FWHM) of 5 mm and high-pass temporal filtering (Gaussian-weighted least squares straight line fitting with sigma = 50 s). Each subject’s native image was normalised to the standard Montreal Neurological Institute (MNI) space via co-registration to their high resolution T1-weighted image. Application of FSL’s MELODIC Independent Components Analysis (ICA; Beckmann & Smith, 2004) separated the fMRI BOLD signal into a set of spatial maps (independent components) representing neural signal and/or noise. An average of 53.17% components were removed across participants’ resting-state fMRI data. Following ICA filtering, low bandpass filtering was applied using AFNI’s ‘*3dBandpass*’ tool (Cox, 1996) to further remove confounding signals below 0.009 Hz and above 0.1 Hz. Prior to analysis, each subject’s corresponding mean functional timeseries image was added back to the bandpass filtered data using *fslmaths* to ensure compatibility with FSL’s FMRI Expert Analysis Tool (FEAT).

*S1.4 Participants, rsfMRI Pre-processing and Analytical Pipeline*

Of the original sample of 96 participants, a total of 77 participants (58 old and 19 young adults) had resting-state fMRI and pulse data. Following quality checks of the rsfMRI and HRV data (including: scanner interference, registration issues, RMSSD values > 200ms), a total of 55 participants (41 old and 14 young adults) were included in the final analyses. The same analytical steps applied to the emotion regulation task-based fMRI data were also performed on the resting-state fMRI data. We conducted both a region of interest functional connectivity analysis to assess the extent to which HRV (obtained during the second session scan) predicted amygdala-mPFC functional connectivity during rest (acquired in the first session). We also examined resting-state whole-brain functional connectivity in the right and left amygdala and the mPFC seeds using FEAT (Woolrich et al., 2004). Clusters surviving a threshold of Z > 3.1 and correction for multiple comparisons with Gaussian random field theory (cluster significance: *p* = 0.05-corrected) were identified (Worsley, 2001).

**Figure S1.** A series of violin plots to display HRV as indexed by (ln)RMSSD values for age group, sex and age group by sex (whole sample, N = 70). **A)** Violin plot displays the significant difference between old (purple, N = 52) and young (light green, N = 18) adults’ HRV as indexed by (ln)RMSSD values (*F*(1,66) = 6.06, *p* = .016, η_p_^2^ = 0.08). Old adults were observed to have significantly lower HRV compared to young adults. **B)** Violin plot displays the (ln)RMSSD values for males (blue) and females (pink) across the whole sample. There was no significant difference in HRV between males and females (*F*(1,66) = 0.09, *p* = .764, η_p_^2^ = 0.00). **C)** Violin plot displays the (ln)RMSSD values for males (blue) and females (pink) per age group. No significant age group by sex interaction for (ln)RMSSD values was found (*F*(1,66) = 0.15, *p* = .698, η_p_^2^ = 0.00). The square represents the mean (In)RMSSD value and the whiskers represent ± 1 standard error around the mean. *(ln)RMSSD*; natural log transformed root mean square of successive differences.

**A)**

**B)**

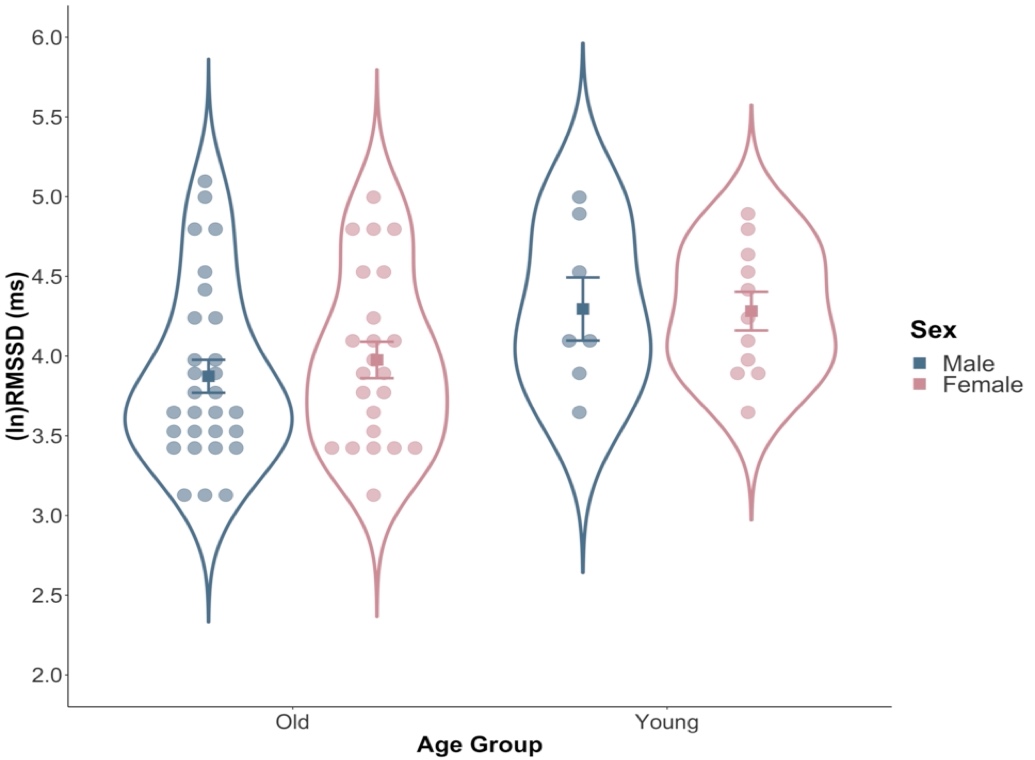

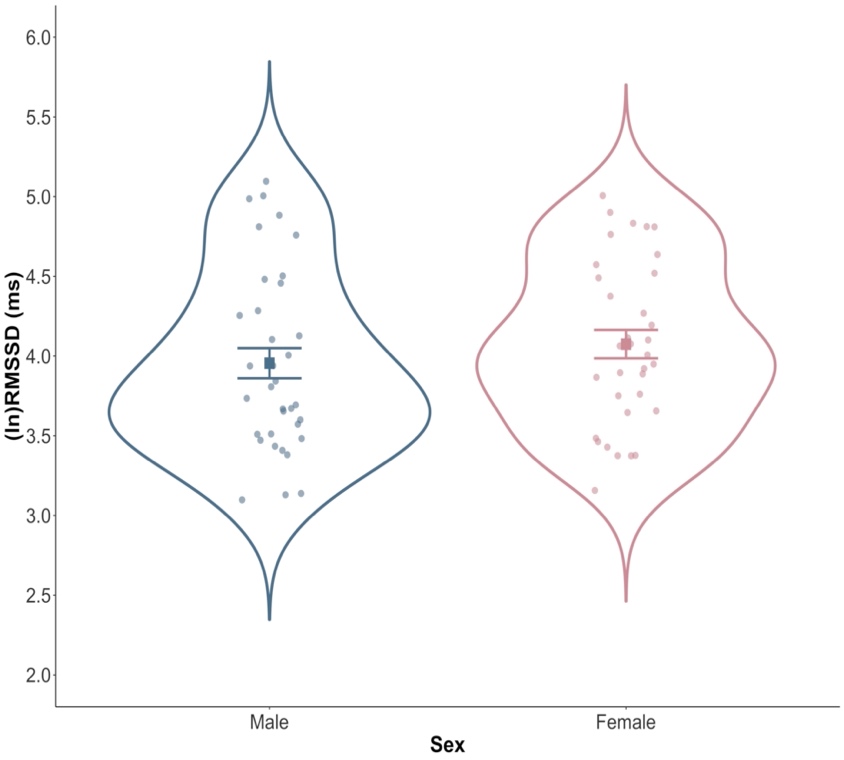

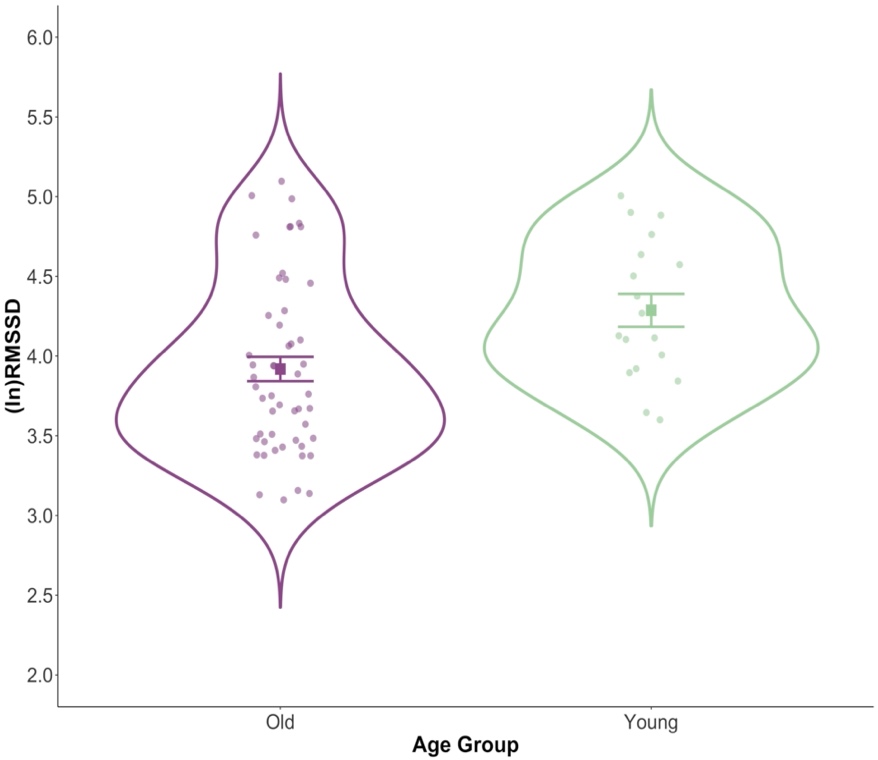

* *p* = .016

**C)**

**B)**

**Figure S2.** The scatterplot displays the natural gap in age (years) between old (purple) and young (light green) adults with natural log transformed RMSSD values ((ln)RMSSD) presented on the y axis.

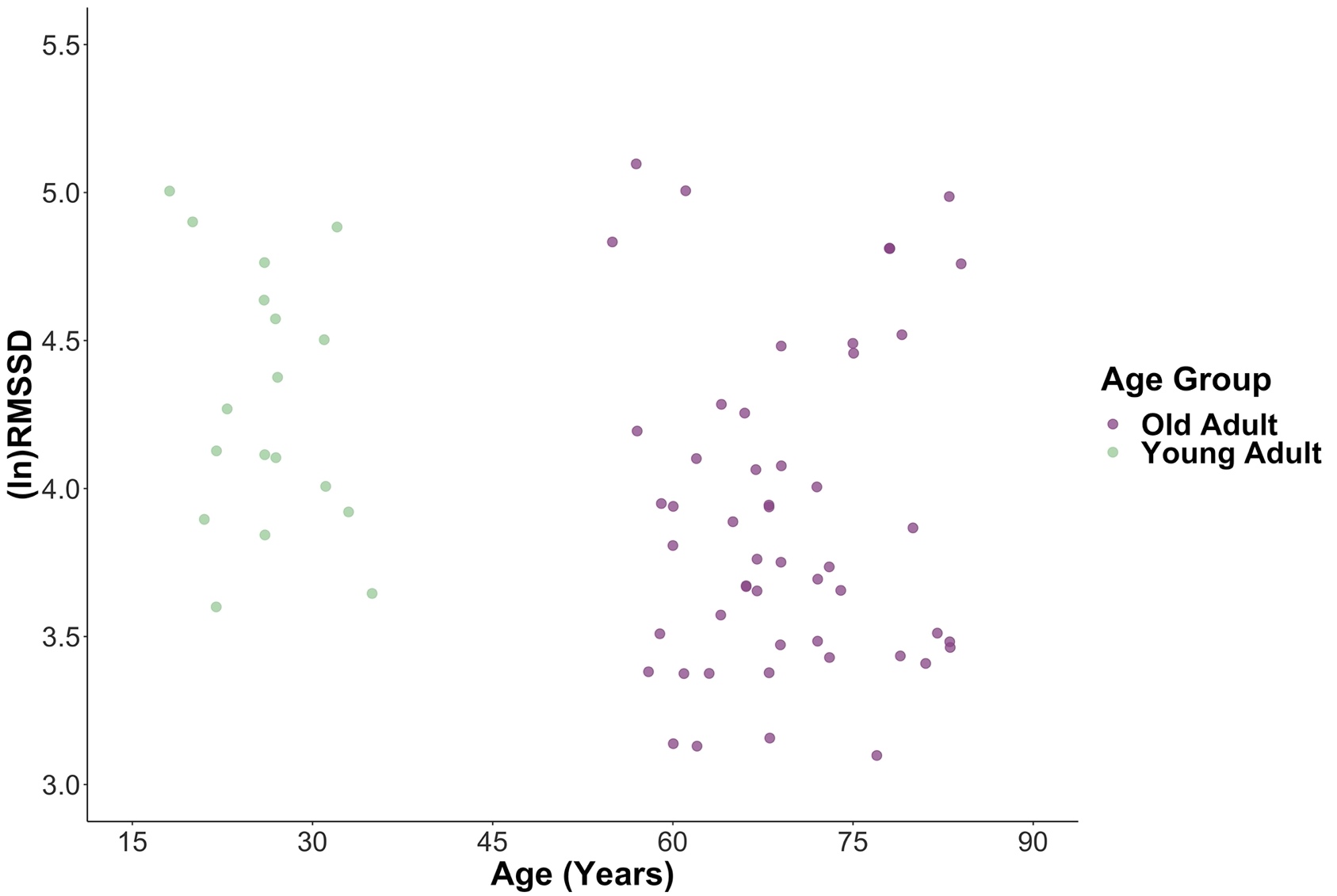

**S2. HRV and Resting-state Amygdala-mPFC Functional Connectivity Analyses**

Multiple regression analyses were employed to examine associations between HRV and resting-state amygdala-mPFC functional connectivity strength in the whole sample (N = 55). Separate multiple regression models were tested with (i) right amygdala-mPFC connectivity and (ii) left amygdala-mPFC connectivity values as dependent variables. The following predictors were entered into the regression model: age group (1 = old adults, 0 = young adults), (ln)RMSSD (centered), and a HRV x age interaction term. In each hierarchical regression model, age group and HRV were entered first (step 1), followed by the HRV x age interaction predictor (step 2) using the enter method. Standardised beta coefficients are reported for all predictors.

*HRV and Resting-state Right Amygdala-mPFC Functional Connectivity*

At step 1, neither the main effect of age (*β* = 0.19, *t* = 1.30, *p* = .198) or HRV (*β* = 0.08, *t* = 0.53, *p* = .596) were found to be significant predictors of the regression model, explaining only 3.20% of the variance in resting-state right amygdala-mPFC functional connectivity (*F*(2,52) = 0.85, *p* = .431). Entering the HRV x age interaction term did not significantly improve the proportion of variance explained in right amygdala-mPFC functional connectivity (Δ*R*^2^ = 0.01) and the overall regression model remained non-significant (*F*(3,51) = 0.80, *p* = .498). Thus, the interaction between HRV and age (*β* = 0.26, *t* = 0.84, *p* = .404) was not found to significantly predict right amygdala-mPFC resting-state functional connectivity.

*HRV and Resting-state Left Amygdala-mPFC Functional Connectivity*

Similar to the right amygdala-mPFC resting-state functional connectivity findings, neither the main effect of age (*β* = -0.11, *t* = -0.74, *p* = .463) or HRV (*β* = 0.10, *t* = 0.69, *p* = .491) were found to be significant predictors of the overall regression model and only explained 2.90% of the variance in resting-state left amygdala-mPFC functional connectivity (*F*(2,52) = 0.78, *p* = .462). When the HRV x age interaction term was subsequently entered into the model, this did not significantly improve the proportion of variance explained in left amygdala-mPFC resting-state connectivity (Δ*R*^2^ = .001) and the overall regression model remained non-significant (*F*(3,51) = 0.52, *p* = .668). Therefore, the interaction between HRV and age (*β* = 0.06, *t* = 0.18, *p* = .855) was not found to significantly predict left amygdala-mPFC resting-state functional connectivity.

| **Table S1. Neural Regions and Local Maxima for Right Amygdala Whole-Brain Resting-state Functional Connectivity as a Function of HRV** | | | | | | | |
| --- | --- | --- | --- | --- | --- | --- | --- |
|  |  |  |  | **MNI Coordinates** | | |  |
| **Region** | **H** | **Cluster Size** | **BA** | **x** | **y** | **z** | **Z** |
| HRV + *(old and young adults)*  **No significant results** |  |  |  |  |  |  |  |
| HRV - (*old and young adults*) |  |  |  |  |  |  |  |
| Precentral Gyrus extending into Postcentral Gyrus | **R** | **83** | **4** | **8** | **-26** | **66** | **4.19** |
|  | R |  | 4 | 12 | -28 | 68 | 4.13 |
|  | R |  | 6 | 8 | -22 | 64 | 3.96 |
|  | R |  | 4 | 14 | -34 | 64 | 3.52 |
| *HRV x Age Interaction + (old and young adults)* |  |  |  |  |  |  |  |
| **No significant results** |  |  |  |  |  |  |  |
| *HRV x Age Interaction - (old and young adults)*  **No significant results** |  |  |  |  |  |  |  |
| HRV + *(old adults)*  **No significant results** |  |  |  |  |  |  |  |
| HRV - *(old adults)*  **No significant results** |  |  |  |  |  |  |  |

Neural regions that demonstrated associations with right amygdala during rest as a function of HRV across the whole sample (N = 55) and old adult sample only (N = 41), Z = 3.1; cluster significance: *p* < 0.05, corrected). Local maxima are listed for clusters containing more than one peak. Cluster size refers to the number of voxels contained within a specific cluster. Coordinates (MNI space) represent location of clusters and their maximum Z-scores (bold) and the location of local maxima within significant clusters and their associated Z-statistic. The Harvard Oxford Structural Cortical and Subcortical atlases within FSL were used to label significant clusters. BA refers to the Brodmann Area for each cluster. The ‘R’ package *label4MRI* (v1.2) was used to generate the BA label based on the MNI coordinates. H = hemisphere (L = left, R = right).

| **Table S2. Neural Regions and Local Maxima for Left Amygdala Whole-Brain Resting-state Functional Connectivity as a Function of HRV** | | | | | | | |
| --- | --- | --- | --- | --- | --- | --- | --- |
|  |  |  |  | **MNI Coordinates** | | |  |
| **Region** | **H** | **Cluster Size** | **BA** | **x** | **y** | **z** | **Z** |
| HRV + *(old and young adults)*  **No significant results** |  |  |  |  |  |  |  |
| HRV - (*old and young adults*)  **No significant results** |  |  |  |  |  |  |  |
| *HRV x Age Interaction + (old and young adults)* |  |  |  |  |  |  |  |
| **No significant results** |  |  |  |  |  |  |  |
| *HRV x Age Interaction - (old and young adults)*  **No significant results** |  |  |  |  |  |  |  |
| HRV + *(old adults)* |  |  |  |  |  |  |  |
| Lingual Gyrus | **R** | **80** | **19** | **12** | **-42** | **-10** | **4.17** |
|  | R |  | 19 | 10 | -50 | -4 | 3.97 |
|  | R |  | 19 | 12 | -52 | -8 | 3.68 |
|  | R |  | 19 | 20 | -46 | -8 | 3.43 |
| HRV - *(old adults)*  **No significant results** |  |  |  |  |  |  |  |
| Neural regions that demonstrated associations with left amygdala during rest as a function of HRV across the whole sample (N = 55) and old adult sample only (N = 41), Z = 3.1; cluster significance: *p* < 0.05, corrected). Local maxima are listed for clusters containing more than one peak. Cluster size refers to the number of voxels contained within a specific cluster. Coordinates (MNI space) represent location of clusters and their maximum Z-scores (bold) and the location of local maxima within significant clusters and their associated Z-statistic. The Harvard Oxford Structural Cortical and Subcortical atlases within FSL were used to label significant clusters. BA refers to the Brodmann Area for each cluster. The ‘R’ package *label4MRI* (v1.2) was used to generate the BA label based on the MNI coordinates. H = hemisphere (L = left, R = right). | | | | | | | |

| **Table S3. Neural Regions and Local Maxima for MPFC Whole-Brain Resting-state Functional Connectivity as a Function of HRV** | | | | | | | |
| --- | --- | --- | --- | --- | --- | --- | --- |
|  |  |  |  | **MNI Coordinates** | | |  |
| **Region** | **H** | **Cluster Size** | **BA** | **x** | **y** | **z** | **Z** |
| HRV + *(old and young adults)*  **No significant results** |  |  |  |  |  |  |  |
| HRV - (*old and young adults*)  **No significant results** |  |  |  |  |  |  |  |
| *HRV x Age Interaction + (old and young adults)* |  |  |  |  |  |  |  |
| **No significant results** |  |  |  |  |  |  |  |
| *HRV x Age Interaction - (old and young adults)* |  |  |  |  |  |  |  |
| White matter extending into Superior Lateral Occipital Cortex | **L** | **79** |  | **-32** | **-64** | **20** | **4.43** |
|  | L |  | 39 | -30 | -74 | 26 | 4.20 |
|  | L |  | 39 | -34 | -70 | 28 | 3.90 |
| *HRV + (old adults)*  **No significant results** |  |  |  |  |  |  |  |
| *HRV - (old adults)*  **No significant results** |  |  |  |  |  |  |  |
| Neural regions that demonstrated associations with the mPFC during rest as a function of HRV across the whole sample (N = 55) and old adult sample only (N = 41), Z = 3.1; cluster significance: *p* < 0.05, corrected). Local maxima are listed for clusters containing more than one peak. Cluster size refers to the number of voxels contained within a specific cluster. Coordinates (MNI space) represent location of clusters and their maximum Z-scores (bold) and the location of local maxima within significant clusters and their associated Z-statistic. The Harvard Oxford Structural Cortical and Subcortical atlases within FSL were used to label significant clusters. BA refers to the Brodmann Area for each cluster. The ‘R’ package *label4MRI* (v1.2) was used to generate the BA label based on the MNI coordinates. H = hemisphere (L = left, R = right). | | | | | | | |

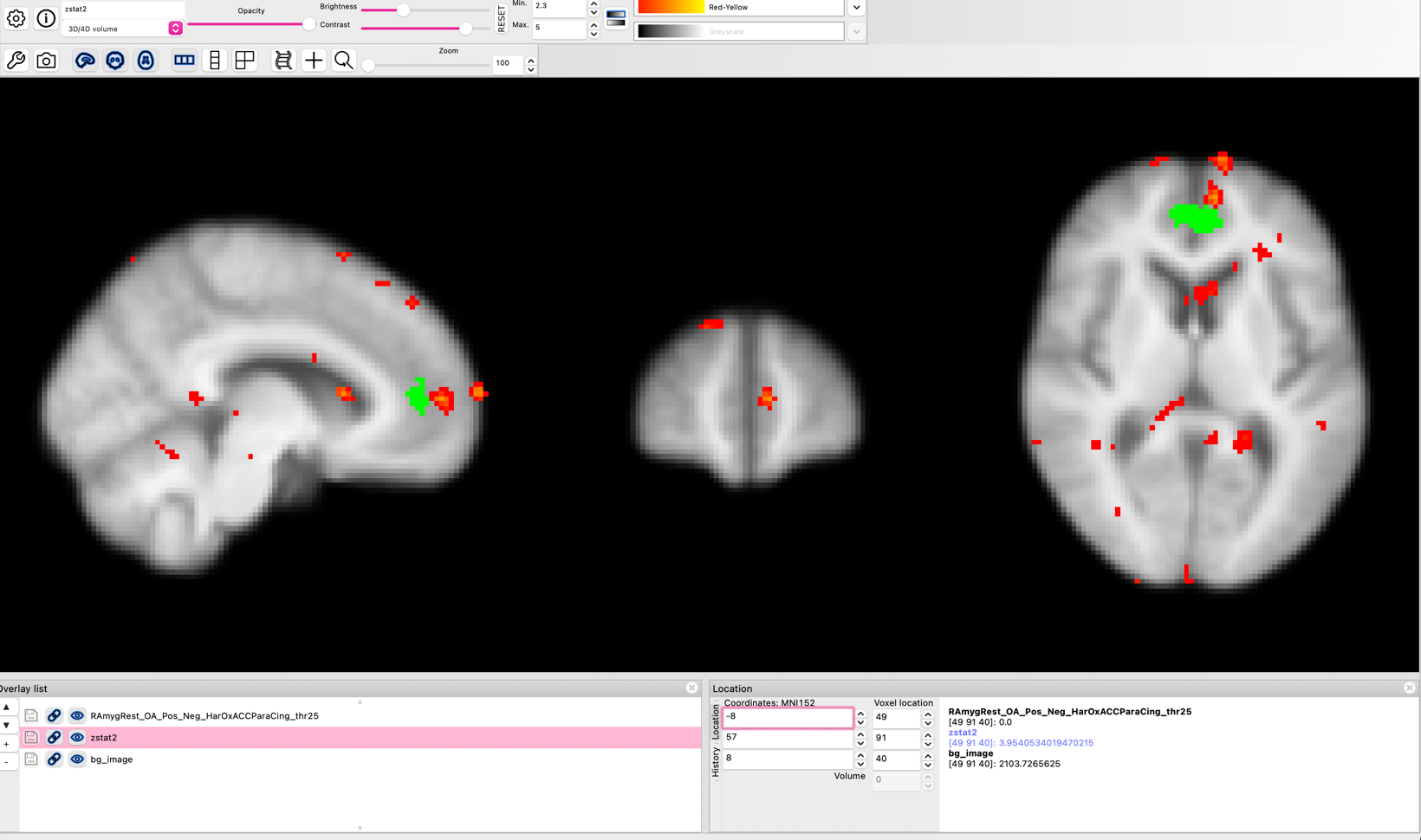

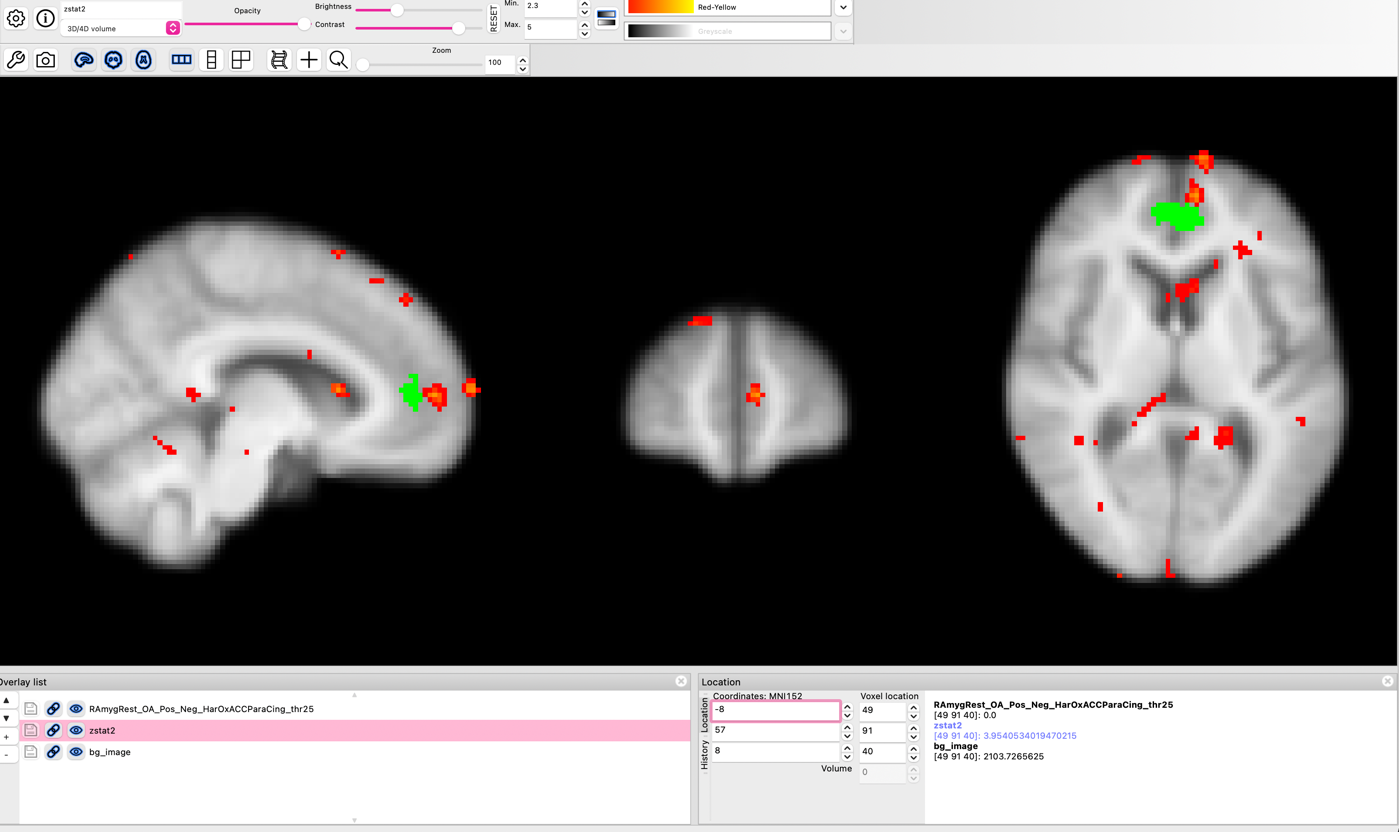

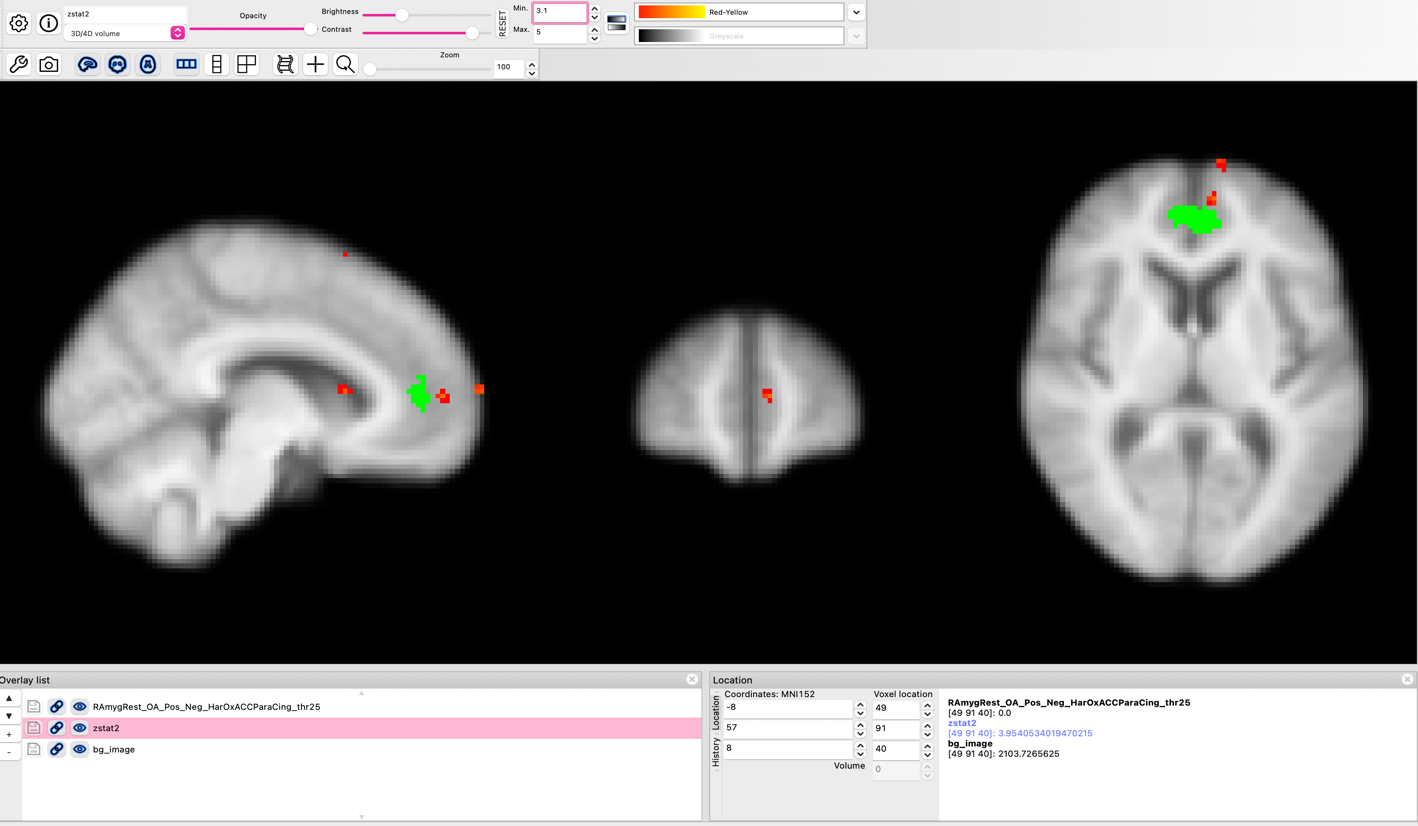

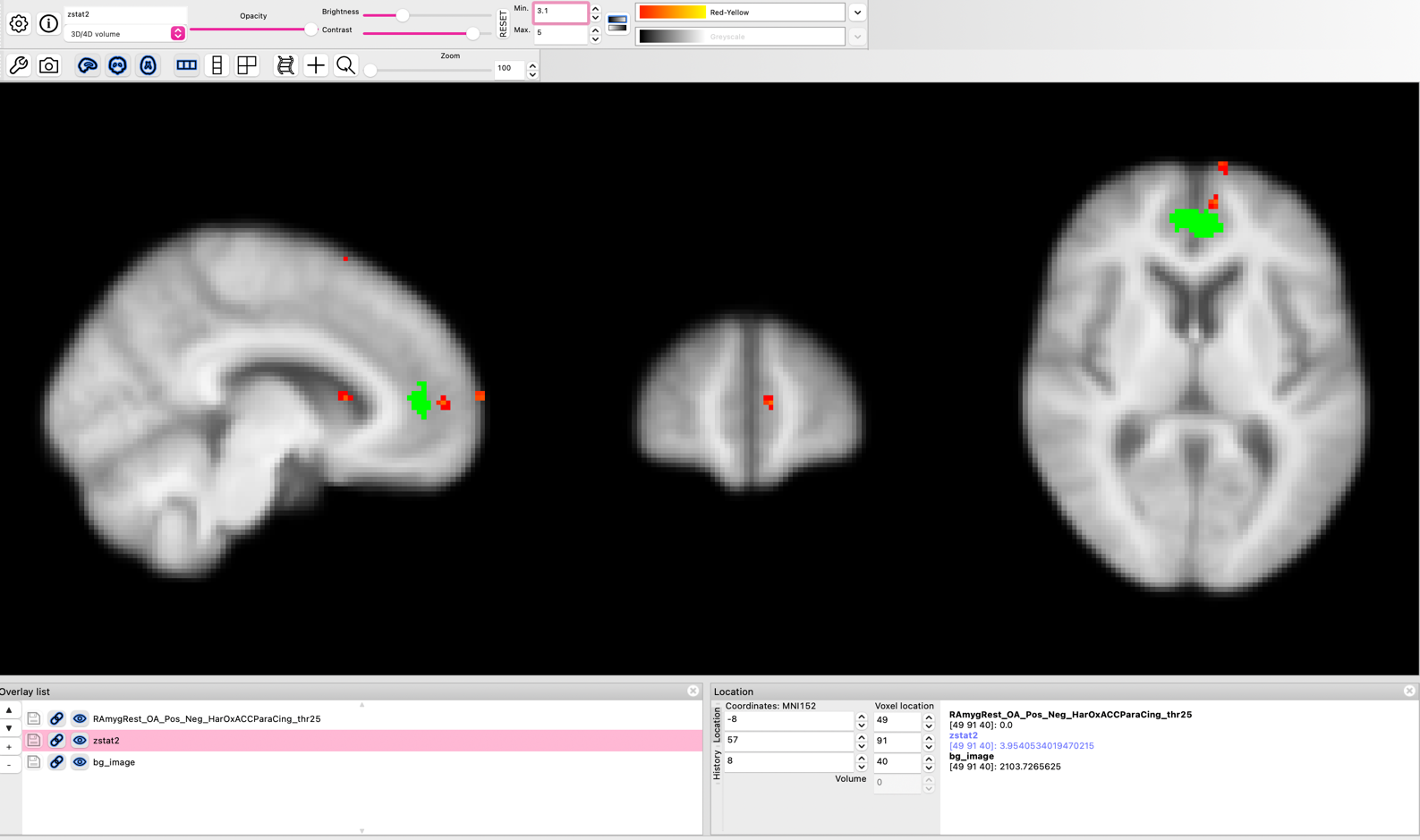

R

R

5.0

2.3

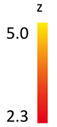

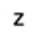

5.0

3.1

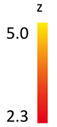

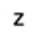

X = -8

Z = 8

**A)**

**B)**

**Figure S3.** Sub-threshold frontal medial cluster which was positively coupled with the left amygdala as a function of higher HRV during rest, but did not survive correction for multiple comparisons (Z = 3.95). The solid green cluster represents the mPFC seed region of interest used in all analyses (Sakaki et al., 2013, 2016). **A)** Frontal medial cluster displayed when Z thresholded at > 2.3. **B)** Frontal medial cluster displayed when Z thresholded at > 3.1.
